# Identification of proteins exhibiting *in vitro* RNA chaperone activity through gradient profiling in the Lyme disease spirochete, *Borrelia burgdorferi*

**DOI:** 10.1101/2025.09.19.677338

**Authors:** Taylor Van Gundy, Cornelius Schneider, Christopher C. Ebmeier, Gavin Chambers, Nick Cramer, Kathryn Flinn, Dan Drecktrah, Jenny A. Hyde, Jon T. Skare, Richard T. Marconi, D. Scott Samuels, Meghan C. Lybecker

## Abstract

RNA-binding proteins (RBPs) play key roles in regulating gene expression in bacteria. However, relatively few RBPs have been discovered and characterized in *Borrelia* (*Borreliella*) *burgdorferi*, the causative agent of Lyme disease. We utilized gradient profiling to identify putative RBPs that co-sediment with small RNAs (sRNAs) and nascent mRNAs. We employed *in vitro* and *in vivo* assays to characterize the RNA chaperone activities of several proteins we identified in the gradient profiling. The previously hypothetical proteins BB0749, BB0713 and BB0796, as well as the chemotaxis-related protein CheY2 and the flagella-associated protein FlgV displayed RNA annealing and/or strand displacement activity. Moreover, *in vivo* Co-IP assays demonstrated BB0749 binds RNA in *B. burgdorferi*.

## Introduction

RNA-binding proteins (RBPs) are involved in cellular processes essential to bacterial survival and fitness; their activities range from enabling translation (ribosome-binding proteins) to regulating transcription, translation, and genome defense. RBPs execute these functions by serving as RNA matchmakers, scaffolds, recruiters, protectors, and structural remodelers (*1, 2*). ‘Conventional’ RBPs bind to sequence and structural motifs of RNA via well-defined RNA-binding domains, such as Sm and Sm-like domains, Fin-O domain, KH domains, double-stranded RNA-binding domains, S1 domain, cold-shock domain, PAZ/PIWI domains, and DEAD box helicase domains (*3, 4*). However, an increasing number of proteins have been identified that lack annotated RNA-binding domains but are capable of binding RNA and, so, have been termed ‘unconventional’, ‘enigma’ or ‘moonlighting’ RNA-binding proteins (*5–10*). Several themes have emerged, including the recurrent identification of metabolic enzymes as RBPs and the frequent association of RNA with proteins containing intrinsically disordered regions (IDRs). Discovery of new and unconventional RBPs has accelerated due to advances in global RBP identification methods utilizing mass spectrometry-based proteomics. Several global RNA-centric approaches have recently been developed for bacterial RNA lacking poly(A) tails. Most of these methods rely on UV crosslinking RNA-protein complexes followed by organic phase extraction or silica-based purification (*8–10*). In contrast, gradient fractionation followed by RNA sequencing (Grad-seq) and its iterations do not require UV crosslinking and rely on separating ribonucleoprotein complexes (RNPs) from cell lysate using a glycerol gradient or size-exclusion chromatography, followed by MS and RNA sequencing on fractions from the gradient (*11–16*). Grad-seq identifies proteins that co-sediment with different classes of RNAs, but not all proteins identified are RNA binding proteins. Gradient sedimentation with RNase treatment (GradR), which includes an RNase treatment to identify RNase sensitive sedimentation profiles, improves the identification of bona fide RNA binding proteins. Grad-seq approaches have been applied to Gram-positive, Gram-negative and cyanobacteria leading to the discovery of novel RBPs and RNA-protein complexes (*17*).

RBPs can transiently interact or form a stable complex with RNA generating RNPs. Stable RNPs in bacteria have diverse functions, ranging from translation via the ribosome, to regulation and matchmaking involving Hfq, ProQ, and CRISPR complexes (recently reviewed by Gerovac *et al.*(*17*)). Stable RNP complexes are characterized by longer lifetimes, lower off-rates (kinetic stability), and higher affinities, with sub-micromolar disassociation constants (thermodynamic stability), compared to transient RNA-RBP interactions. Transient interactions between RNA and RBPs are inherently short-lived, enabling the RBPs to dynamically regulate diverse processes in response to cellular signals. Many mechanisms of RNA-based gene regulation rely on RNA-protein interactions to modulate RNA structures, facilitate RNA-RNA interactions, or occlude binding sites of other proteins or RNAs (*1–3, 18*). RNA chaperones are a group of RBPs capable of disrupting RNA secondary and tertiary structure (unwinding, unfolding, and strand displacing) or accelerating base-pairing of RNAs (annealing). RNA chaperones perform their functions through both stable and/or transient interactions with RNA (*19–24*).

*Borrelia (Borreliella) burgdorferi* is the causative agent of Lyme disease and casts a wide shadow in the United States with an estimated 476,000 patients treated annually (*25*). *B. burgdorferi* cycles in nature between a tick vector and vertebrate host, requiring phase-specific gene regulation. Transcriptomic studies have generated comprehensive maps of differentially regulated genes under various biologically relevant growth conditions *in vitro* and, more recently, the transcriptome has also been defined *in vivo* during tick feeding (*26–28*). However, our understanding of posttranscriptional gene regulation remains limited. RBPs posttranscriptionally regulate gene expression by interacting with many, if not most, RNAs, including rRNAs, tRNAs, mRNAs, and regulatory RNAs such as *trans*-encoded small RNAs (sRNAs) and antisense RNAs (asRNAs). A large number of sRNAs have been identified in *B. burgdorferi* via transcriptomic studies (*29–34*), but only a few have been characterized (*35, 36*). A handful of *B. burgdorferi* proteins have been reported to have RNA-binding capabilities, including BpuR, BosR, EbfC, SpoVG, CsrA, KhpA, KhpB, PlzA, and FlgV (previously annotated as HfqBb) (*26, 37–46*). However, the RNA chaperone activity, mechanisms of regulation and comprehensive RNA interactomes have not been identified for these proteins.

In this study, we employed gradient fractionation of RNPs followed by mass spectrometry to identify putative RNA binding proteins and characterize the RNA chaperone activity of a selected subset of them. We also chose to examine the RNA chaperone activity of a handful of proteins that were annotated or reported as nucleic acid-binding proteins. Most of the nucleic acid-binding proteins demonstrated RNA chaperone activity, except for the single-stranded DNA binding protein, SSB. The four putative RBPs (BB0749, BB0713, BB0796, and CheY2) also demonstrated strong RNA chaperone activity. Co-immunoprecipitation (Co-IP) assays show that BB0749 binds RNA in *B. burgdorferi*. Future studies will focus on the identification of binding partners and mechanisms of the newly identified RBPs.

## Results and discussion

### Glycerol gradient fractionation of B. burgdorferi RNA-protein complexes

To identify potential RBPs and RNA chaperones, we fractionated *B. burgdorferi* cell lysate on a glycerol gradient to separate RNP complexes by sedimentation as seminally described by Smirnov *et al.* (*11*) in gradient sequencing (Grad-seq) (Fig. 1A). Following sedimentation, 20 fractions were collected, and each fraction was analyzed for both RNA and protein constituents on denaturing gels (Fig. S1). The fractionation pattern of RNAs and proteins in the gradient demonstrates successful sedimentation and separation of RNP complexes. sRNAs, including tRNAs, were detected in fractions 1-5, while rRNA was detected in fractions 9-18 (Fig. S1). In addition, RNA polymerase (RNAP) subunits and 6S RNA fractionated together as expected primarily in fractions 4-7, and rRNA and 5S RNA fractionated together in fractions 9-18, consistent with what was reported previously in other organisms (Fig. 1B and S1) (*11–17*). In Grad-seq and its iterations, the RNA and proteins found in each Grad-seq fraction were identified by RNA-seq and mass spectrometry, respectively (*11, 15, 16*). However, we were particularly interested in identifying proteins that bind sRNAs and nascent mRNAs and proceeded with LC MS/MS of fractions 1-10. One hundred and forty-five proteins were identified in fractions 1-10 and were manually classified by annotated and predicted functions (Fig. S2 and Table S1). The fractionation pattern of a subset of these proteins identified by LC MS/MS, including annotated or reported RNA and/or DNA-binding proteins (SSB, RRF, BosR, BpuR, SpoVG, KhpA, KhpB, and EbfC) are depicted in Fig. 1B. Most of the recognized nucleic acid-binding proteins, BosR, BpuR, EbfC, RRF, and SSB, along with uncharacterized proteins BB0749 and BB0796, are found predominantly in fractions 1-5, which co-sediment with sRNAs. Uncharacterized protein BB0713, KhpB and SpoVG resolved in fractions 4-8 together with the RNAP subunits, while KhpA was identified in all fractions, but was more abundant in fractions 3-8. Western blot analyses of BosR, BB0749, and RNAP β-subunit in fractions 1-10 validate mass spectrometry results (Fig. 1B).

**Figure 1.**
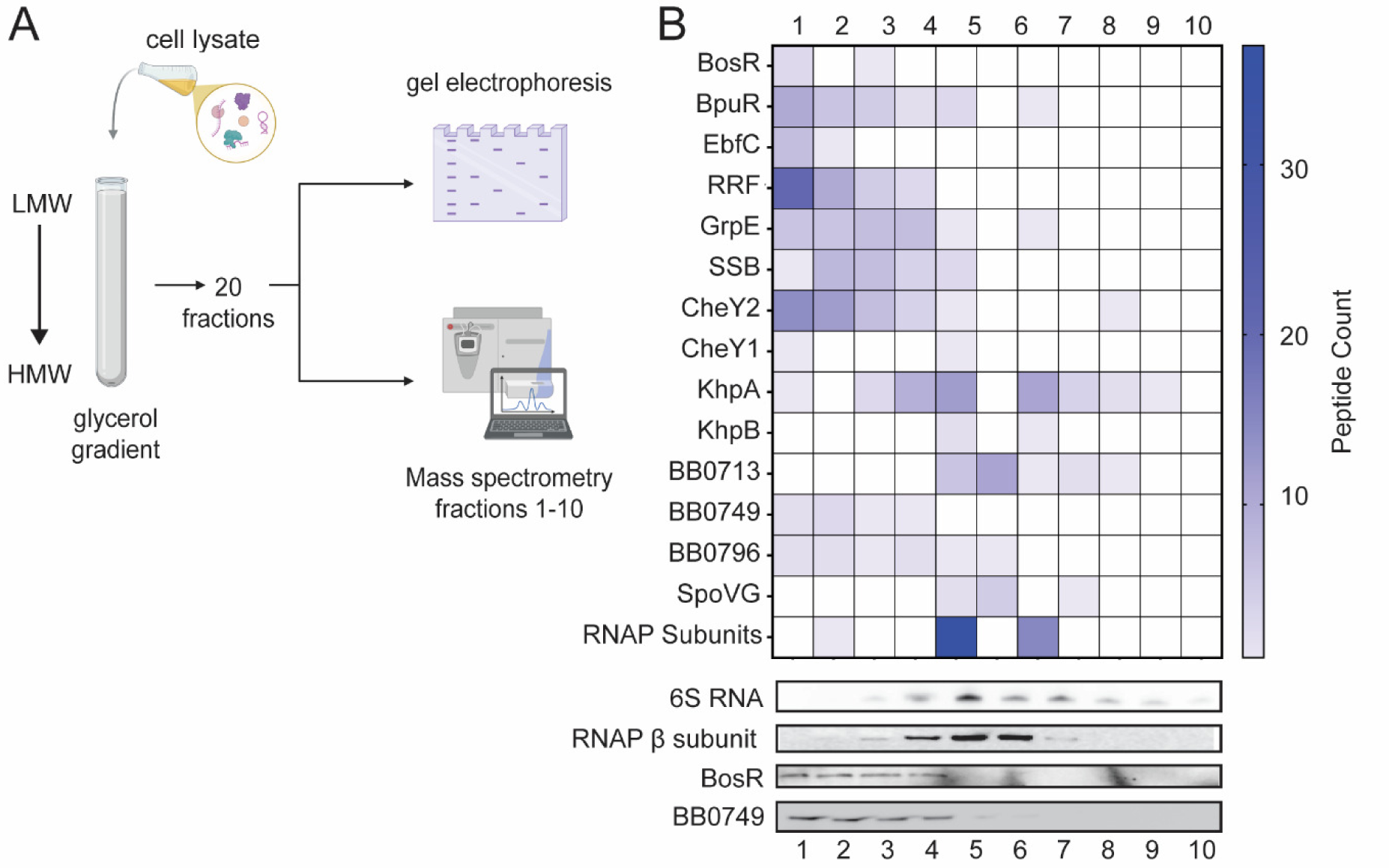
Glycerol gradient fractionating *B. burgdorferi* RNA-protein complexes. (A) Schematic of glycerol gradient experimental strategy. *B. burgdorferi* cell lysate was fractionated in a glycerol gradient. Low molecular weight (LMW) complexes resolve toward the top of the gradient and high molecular weight (HMW) complexes resolve toward the bottom. 20 fractions were collected and analyzed for protein and RNA content. Mass spectrometry was performed on fractions 1-10. (B) Heat map of a subset of proteins identified in fractions 1-10. A 6S RNA Northern blot and RNAP β-subunit Western blot demonstrates successful fractionation of RNPs and validate MS analyses. Western blots of BosR and BB0749 also validate MS gradient data.

Not all proteins that co-sediment with specific classes of RNAs are RBPs or components of RNA complexes. Proteins identified as putative RBPs through Grad-seq are often validated as bona fide RBPs using Co-IPs of RNA with the protein or other orthogonal approaches. Co-IP assays require epitope-tagged proteins or highly specific antibodies, both of which are labor intensive and present certain limitations. Epitope-tagging can alter the native structure and function of the protein and generating antibodies with sufficient specificity is not always successful. Moreover, genetic manipulation of *B. burgdorferi* takes longer than in most bacteria due to its slow growth, further complicating generation of epitope-tagged proteins. Therefore, we choose to biochemically analyze a subset of proteins identified through gradient profiling to assess their *in vitro* RNA chaperone activity before assaying *in vivo* by RNA Co-IP. Selection criteria of proteins included their co-sedimentation with sRNAs and/or RNAP, as well as their predicted structures and/or annotated domains (Table S2). BB0749 is an uncharacterized protein that has a 40-residue IDR. Despite their lack of stable structure, IDRs can function as the sole RNA binding domain of an RBP and can transition to an ordered state upon binding to RNA (*5*). BB0749 co-sedimented in fractions 1-3 but does not contain any annotated domains known to bind RNA, other than an IDR. Uncharacterized protein BB0713 has an annotated *N*-terminal coiled-coil domain and a *C*-terminal C4-type zinc-ribbon domain. The zinc-ribbon domain is rich in aromatic and positively charged amino acids and consists of two Zn knuckles scaffolded by two β-strands. Aromatic and positively charged amino acids are often residues important for RNA binding. In addition, the Zn knuckle is a zinc finger subtype known to interact with RNA (*4*). A *Helicobacter pylori* protein with the same Zn ribbon 9 domain binds *flaA* mRNA (*47*). CheY2 is annotated as a chemotaxis response regulator and is necessary for vertebrate infection (*48*). The *B. burgdorferi* genome encodes three CheY response regulators (CheY1-3) but only CheY3 was found to be essential for motility and chemotaxis (*49*). Xu *et al.* (*48*) hypothesized that CheY2 functions as a transcriptional or posttranscriptional regulator of virulence genes. None of the CheY response regulators in *B. burgdorferi* contain annotated RNA-binding or DNA-binding motifs. We also chose to examine the RNA chaperone activity of several known or predicted nucleic acid-binding proteins (BosR, BpuR, FlgV, KhpA, EbfC, RRF, and SSB).

### RNA annealing and strand displacement activities of putative RBPs

RNA chaperones are RBPs that restructure RNA by disrupting secondary and tertiary structures (unwinding, unfolding, and/or strand displacement) or accelerate base-pairing (annealing). We employed an in-solution coupled RNA annealing and strand displacement assay to determine RNA chaperone activities of a subset of the putative RBPs identified in gradient profiling. The assay was originally developed by the Schroeder lab and utilized fluorescence resonance energy transfer (FRET) to measure the annealing and strand displacement rates of RNAs labeled with Cy3 and Cy5 (*21, 50–54*). We modified the assay to measure a fluorescent signal from a fluorophore that is modulated by a quencher, rather than FRET (Fig. 2A). Two previously published fully complimentary 21-mer unstructured RNAs were used in the assay (*46, 54*). M1 RNA was 5′ labeled with a 6-FAM fluorophore (FAM-M1) and J1 was 3′ labeled with Dabcyl (J1-Dab) (Table S3). RNA annealing (phase I) was initiated by injecting J1-Dab into a reaction mixture containing equimolar amounts of FAM-M1, either in the presence or absence of protein at 37°C. The fluorescent signal was quenched over time as J1-Dab RNA base-paired with FAM-M1. Strand displacement (phase II) was initiated by injection of tenfold excess unlabeled M1 (uM1) into the reaction and the fluorescent signal increased over time as uM1 replaced FAM-M1 in the duplex with J1-Dab, releasing the fluorophore from the quencher.

**Figure 2.**
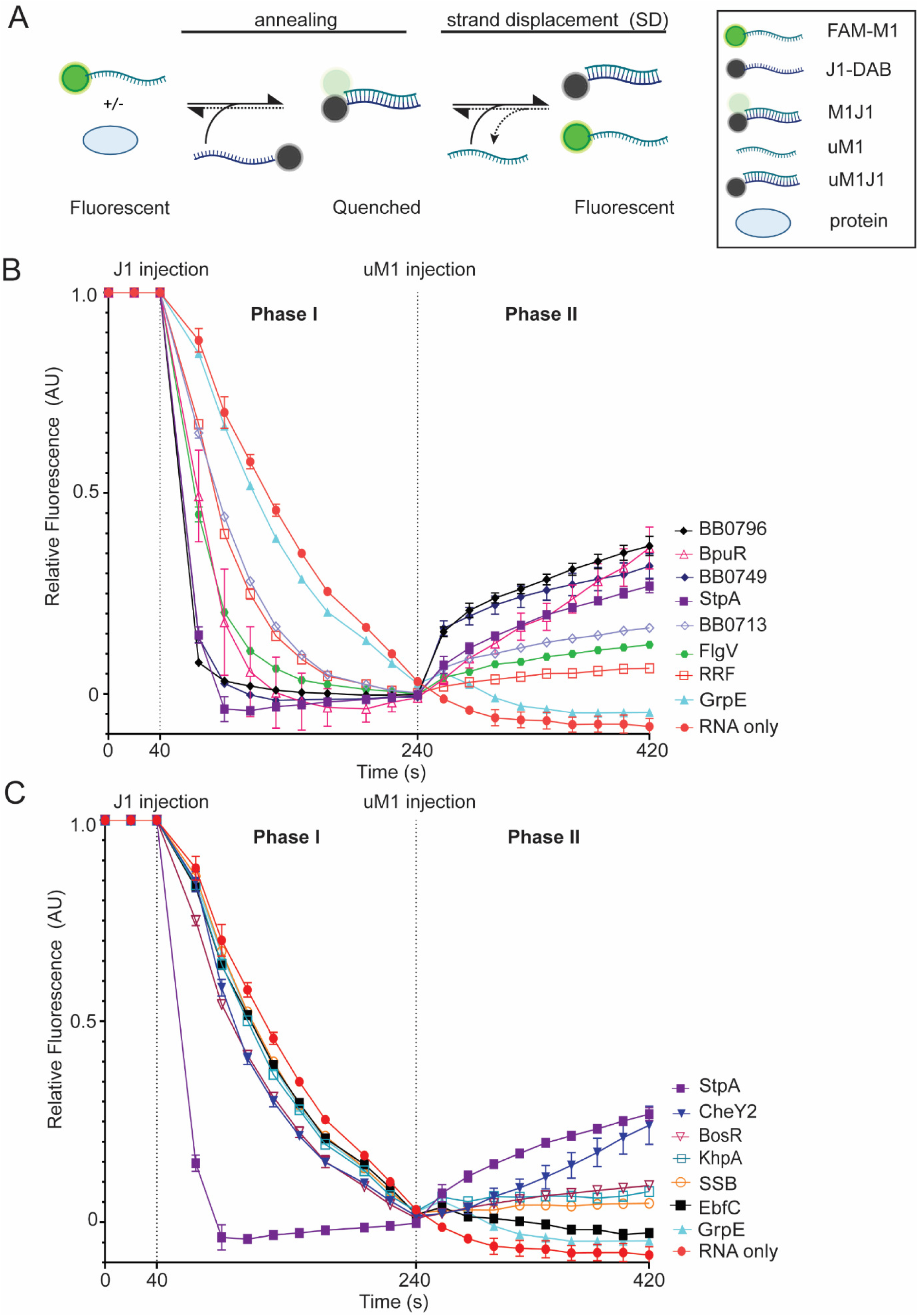
*B. burgdorferi* proteins with in-solution RNA annealing and strand displacement assay. (A) Schematic of in-solution annealing and strand displacement experimental strategy. The fluorescence of the 5′ FAM-labeled M1 RNA (FAM-M1), with or without protein, is measured for 40 s. The RNA annealing reaction is started by injection of 3′ Dabcyl-labeled J1 (J1-Dab) RNA. Fluorescent measurements were taken every 10 s for 200 s. Base-pairing of FAM-M1 and J1-Dab results in quenching of the FAM-fluorophore (M1J1). Unlabeled-M1 (uM1) is injected at 240 s and fluorescence is measured for another 200 s, with 10 s intervals. Strand displacement is observed as uM1 replaces FAM-M1 in the duplex (uM1J1), releasing FAM-M1 and increasing fluorescence. (B and C) Time-resolved curves were normalized to 1 at 40 s and 0 at 240 s. Each vertical dotted line represents the time that RNA was injected into each sample. RNA without protein (red circle) or GrpE (teal triangle) served as negative controls. StpA, an RNA chaperone from *E. coli*, was used as a positive control (purple square). *B. burgdorferi* proteins BpuR (open pink triangle), FlgV (green hexagon), RRF (open red square), BB0713 (open lavender diamond), BB0749 (blue diamond) and BB0796 (black diamond) with annealing rates at least 2-fold higher than RNA only are displayed in (B). Whereas *B. burgdorferi* proteins BosR (open magenta upside-down triangle), CheY2 (blue upside-down triangle), EbfC (black square), KhpA (open teal square) and SSB (open orange circle) with annealing rates less than 2-fold compared to RNA only are displayed in (C). Error bars represent the standard error of the mean (SEM) of three replicates.

The fluorescent signal of the reaction at the start of the assay is variable depending on the protein added to the reaction (Fig. S3A and B). Therefore, this assay focuses on the kinetics of the reactions. To compare and visualize the results the fluorescent signal was normalized to 0 at the time point (240 s) directly before uM1 injection (Fig. S3C and D) and normalized to 1 at the time point (40 s) direct prior to J1-Dab injection (Fig. S3E and F). In the absence of protein (RNA only), the short fully complementary unstructured RNAs used in the assay anneal with an observed annealing rate constant of 3.15 × 10^-6^ s^-1^ (Fig. 2 and Table S4). *E. coli* StpA, an RNA chaperone with known annealing and strand displacement activity, was used as a positive control and *B. burgdorferi* GrpE was used as a negative control, as previously described (*46*). As expected, StpA accelerated the rate of RNA annealing 17.3-fold compared to RNA alone, while GrpE demonstrated minimal to no RNA annealing compared to RNA alone (Fig. 2B and C). *B. burgdorferi* BpuR, FlgV, RRF, BB0713, BB0749, and BB0796 proteins accelerated the rate of RNA annealing by over twofold compared to RNA only (5.4-fold, 5.4-fold, 2.7-fold, 2.5-fold, 16.5-fold, and 21.7-fold) (Fig. 2B and Table S4). While BosR, CheY2, KhpA, SSB and EbfC accelerated RNA annealing by less than twofold compared to RNA alone (1.5-fold, 1.6-fold, 1.2-fold, 1.2-fold, and 1.1-fold) (Fig. 2C and Table S4).

Strand displacement (phase II) of the reaction is initiated by the injection of tenfold molar excess of uM1 (Fig 2). If strand displacement occurs, then uM1 replaces FAM-M1 in the duplex with J1-Dab, resulting in increased in fluorescence. However, simultaneous annealing of remaining single-stranded J1-Dab with FAM-M1 or uM1 can continue, resulting in transient fluorescent quenching until the remainder of J1-dab is fully annealed. The extent and detection of annealing during strand displacement depends on the amount of single-stranded FAM-M1 at the initiation of strand displacement as well as the presence of proteins with strand displacement and/or annealing activity. For instance, in a reaction with a protein that has both annealing and strand displacement activity (StpA), FAM-M1 is largely duplexed with J1-Dab at the end of phase I, and the fluorescence is quenched and plateaued (Fig. 2 and S3). Upon uM1 injection, strand displacement is initiated and FAM-M1 is replaced in the duplex with uM1 and fluorescence increases. Any remaining FAM-M1 and J1-Dab annealing is not detected during strand displacement reaction (Fig. 2 and S3). In a reaction with a protein with strong annealing activity and minimal strand displacement activity (e.g., RRF), fluorescence is quenched and plateaued at the end of phase I. Again, most of FAM-M1 is duplexed with J1-Dab and continued annealing is not detected after initiation of phase II. However, the fluorescent signal only increases slightly and remains relatively stable, suggesting little to no strand displacement activity (Fig. 2B and S3). In the absence of protein (RNA only) or a protein lacking annealing or strand displacement activity (e.g., GrpE), RNA annealing continues after initiation of strand displacement. The rate of annealing is much slower in these reactions and more single-stranded FAM-M1 and J1-Dab remain at the time of uM1 injection. However, after uM1 injection the ratio of uM1:FAM-M1 is high and most of J1-Dab anneals with uM1, leading to reduced quenching and a slower apparent annealing rate, compared to phase I. Consequently, we cannot directly compare the strand displacement and negative control’s reaction rates. Therefore, a protein is considered to have strand displacement activity in this assay if the strand displacement rate is positive (increasing fluorescence), and the fluorescence must increase at least 1000 arbitrary units (AU) during phase II in the unnormalized data (Fig. S3, S4 and Table S4). As anticipated, strand displacement does not occur in the RNA only and GrpE reactions, and StpA has robust strand displacement activity. BosR, BpuR, CheY2, FlgV, BB0713, BB0749, and BB0796 all demonstrate strand displacement activity, while RRF, SSB, KhpA, and EbfC do not (Fig. 2, S4 and Table S4).

Dipartite RNA gel annealing and strand displacement assays were also conducted to examine RNA chaperone activity of the proteins as previously described (*46*). Cy3-labeled M1 and Cy5-labeled J1 RNAs were used in the gel assays to observe RNA annealing and strand displacement separately. To start the gel RNA annealing assay, Cy5-labeled J1 was added to a master mix containing Cy3-labeled M1 in the presence or absence of a protein. Samples were removed from the reaction over time, mixed with stop buffer containing an excess of unlabeled M1 RNA (uM1) and resolved by electrophoresis on a native polyacrylamide gel (Fig. 3A). RNA annealing is detected as the formation of double-stranded J1M1 RNA (yellow band), representative images for each protein are shown in Fig. S5A. The formation of J1M1 (RNA annealing) over time was plotted and quantified using the ratio of dsRNA (J1M1) to total RNA (J1M1 and J1uM1) using only the Cy5 signal (Fig. 3B). As expected, StpA increased the rate of RNA annealing by 8.8-fold compared to the RNA only control and the amount of dsRNA formed at the endpoint of the assay was significantly higher than RNA only (Fig. 3B, S5B and Table S4). BosR, BpuR, CheY2, EbfC, FlgV, KhpA, RRF, BB0713, BB0749, and BB0796 all accelerated RNA annealing (5-fold, 2.7-fold, 7.7-fold, 3.0-fold, 18.4-fold, 2.1-fold, 4.7-fold, 4.3-fold, 8.3-fold and 2.1-fold) compared to RNA only and formed significantly more dsRNA at the endpoint of the assay compared to RNA alone. SSB and the negative control GrpE did not demonstrate RNA annealing activity in the gel assay (Fig. 3B, S5B and Table S4). BpuR, FlgV, RRF, BB0713, BB0749, and BB0796 accelerated the rate of annealing by more than twofold compared to RNA only in both the in-solution and gel-based assay. However, BosR, CheY2, EbfC, and KhpA increased the rate of annealing less than twofold compared to RNA alone in the in-solution assay but performed better in the in-gel assay demonstrating greater than twofold increase in annealing rate. These differences may be attributed to the temperature conditions under which the assays were performed (23°C for the gel-based assay and 37°C for the in-solution assay), the difference in the length of time of the assays or to differences in the reaction milieu between the two assays, including the 5’ and 3’ modifications of the RNAs.

**Figure 3.**
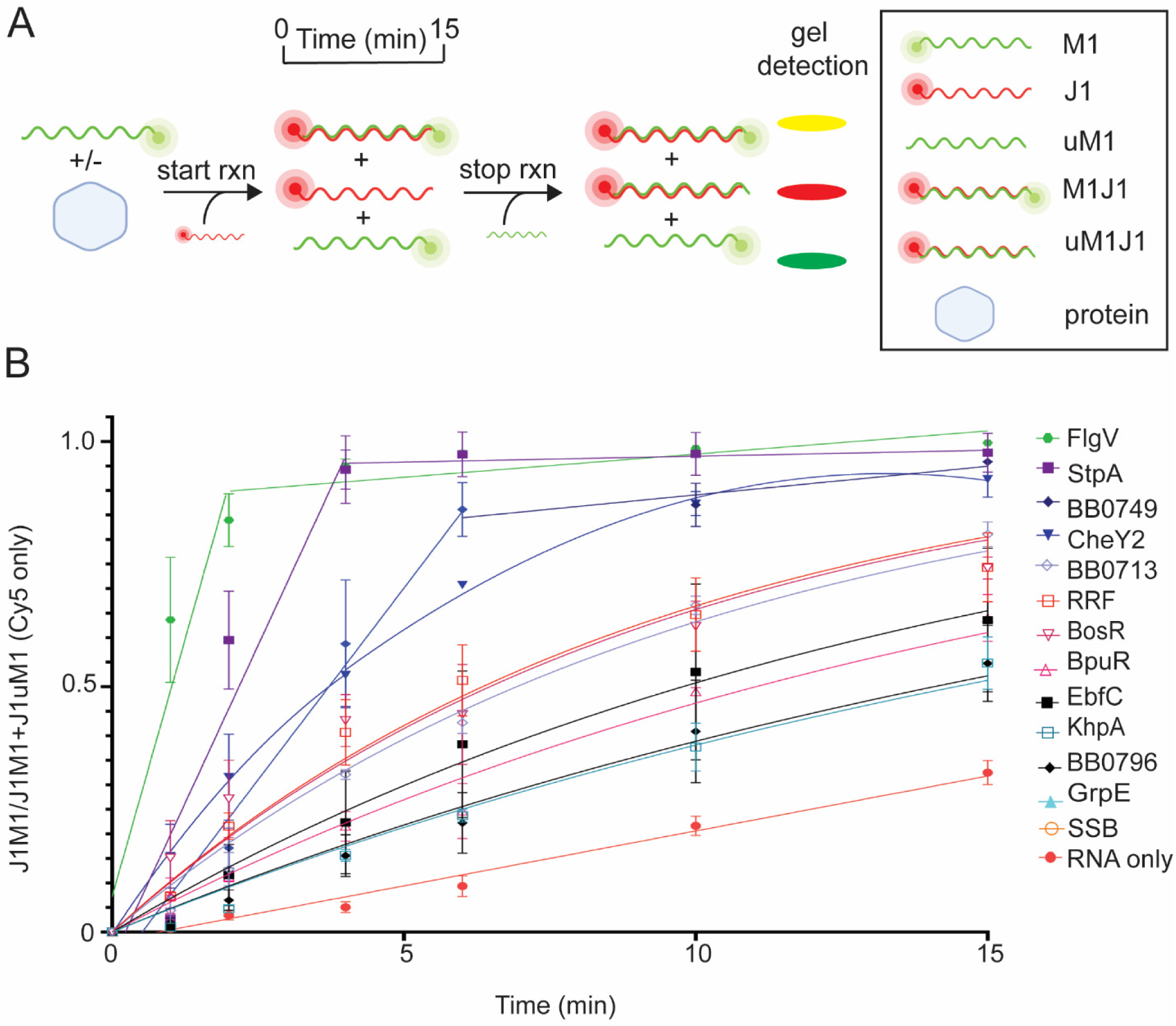
*B. burgdorferi* proteins with gel RNA annealing activity. (A) Schematic of RNA annealing assay using Cy3-labeled M1 (green) and Cy5-labeled J1 (red) RNAs. The reaction is initiated by adding Cy5-labeled J1 to a mixture of Cy3-labeled M1 RNA and buffer with or without protein. Annealing is measured over a time course of 15 min resulting in an increasing yellow band (J1M1). Stop buffer and excess unlabeled-M1 (uM1) are used to halt the reaction. Excess uM1 binds to J1, yielding a dsRNA that appears as a red band (J1uM1). This band is slightly smaller than J1M1 due to the lack of the Cy3-fluorophore on uM1. Representative images of RNA annealing gels are shown in Fig. S4A. (B) Normalized linear regression of the ratio of J1M1 dsRNA to total RNA at each timepoint produced the annealing curves for each sample. RNA without protein (red circle) and StpA (purple square) served as negative and positive controls, respectively. FlgV (green hexagon), BB0749 (blue diamond), CheY2 (blue upside-down triangle), BB0713 (open lavender diamond), RRF (open red square), BosR (open magenta upside-down triangle), BpuR (open pink triangle), EbfC (black square), KhpA (open teal square), and BB0796 (black diamond) displayed gel annealing activity. Error bars represent the SEM of three replicates.

Gel strand displacement assays were initiated by adding unlabeled-M1 competitor RNA into a reaction that contained preformed dsRNA (Cy3-labeled M1 and Cy5-labeled J1). Samples were removed from the reaction over time and resolved by electrophoresis on a native polyacrylamide gel (Fig. 4A). Strand displacement was detected by the loss of M1J1 dsRNA (yellow band) and the formation of uM1J1 dsRNA (red band) representative images are shown in Fig. S6. The dissolution of M1J1 (yellow band) and the formation of uM1JI (red dsRNA band) was plotted over time and quantified using the ratio of dsRNA (J1M1) to total RNA (J1M1 and J1uM1) using only the Cy5 signal (Fig. 4B). As noted earlier, strand displacement does not occur spontaneously at 37°C under these conditions and, consistent with this, strand displacement was not observed in the RNA only or GrpE reactions, whereas StpA increased the rate of strand displacement 16.5-fold compared to RNA alone and had significantly less of the dsRNA M1J1 at the end of the assay compared to RNA alone (Fig. 4B, S6C and Table S4). BB0749 also demonstrated strand displacement activity, increasing the rate of strand displacement 20.6-fold compared to RNA alone, while CheY2 and FlgV exhibited modest strand displacement activity, increasing the rate of strand displacement 2.6- and 2.8-fold compared to RNA alone (Fig. 4B and Table S4). BB0749, CheY2 and FlgV also finished the assay with significantly less of the dsRNA M1J1 compared to RNA alone (Fig. S6C). All the other proteins (BpuR, BB0713, BB0796, BosR, KhpA, EbfC, RRF, and SSB) lacked strand displacement activity in the gel assay (Fig. S6B).

**Figure 4.**
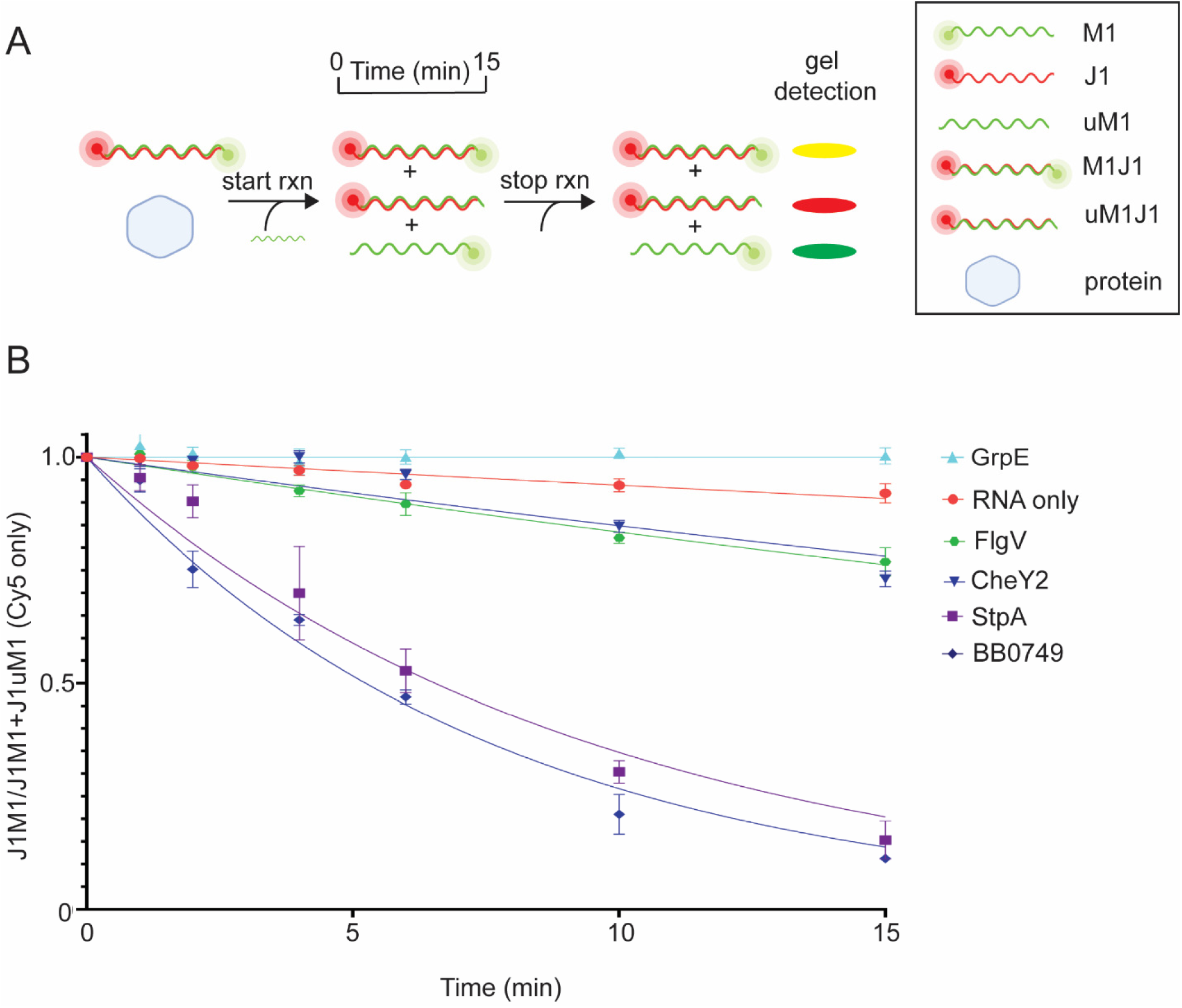
*B. burgdorferi* proteins with gel RNA strand displacement activity. (A) Illustration of the gel RNA displacement assay. Preformed dsRNA (M1J1) was generated with 5′ Cy3-labeled M1 and 5′ Cy5-labeled J1. The strand displacement reaction is initiated by adding unlabeled M1 (uM1) to J1M1 in the presence or absence of protein. Samples are withdrawn from the reaction over time and analyzed by native polyacrylamide gel electrophoresis. The amount of preformed J1M1 dsRNA (yellow band) will decrease with a concomitant increase in the J1uM1 dsRNA (red band) formation over time if the protein has strand displacement activity. Representative images of the RNA strand displacement gels are shown in Figure S5A. (B) Normalized non-linear regression of the ratio of J1M1 dsRNA to total RNA for each time point yielded stand displacement curves. Only the Cy5 channel was quantified and used when graphing the curves. RNA only (red circle) and StpA (purple square) were used in the assay as negative and positive controls, respectively. FlgV (green hexagon), BB0749 (blue diamond), and CheY2 (blue upside-down triangle) displayed gel strand displacement activity. Error bars represent the SEM of three replicates.

KhpA, EbfC, RRF and SSB did not demonstrate strand displacement activity in either the in-solution or gel-based assay. While BB0713, BB0796, BosR, and BpuR demonstrated strand displacement activity in the in-solution but not gel-based strand displacement assay (Table S4). Strand displacement of one RNA in a duplex by another RNA is a concerted process of opening the initial double strand and annealing of the new duplex. RBPs with strand displacement activity destabilize double strands, either from the ends or bulges in base-paired regions. A third strand uses the destabilized regions to initiate invasion. RBPs that strand displace and exhibit RNA annealing activity can catalyze strand displacement by both destabilizing double stranded regions and accelerating the annealing reaction of the invading strand. The in-solution and gel-based assays use the same RNA sequence, but have different fluorophores conjugated to different ends of the RNA, which may explain the different results between the two assays for some proteins. The in-solution assay has the 6-FAM fluorophore on the 5ʹ end of the M1 RNA and the quencher on the 3ʹ end of the J1 RNA, while the gel-based assay has Cy3 and Cy5 fluorophores attached to the 5ʹ ends of M1 and J1. Cy3 and Cy5 dyes linked to the 5ʹ ends of a DNA duplex have been reported to stabilize the duplex (*55, 56*), and this may explain why some proteins did not display strand displacement activity in the gel-based assay. Moreover, having the 6-FAM and Dabcyl quencher proximal to each other on the RNA duplex in the in-solution assay may destabilize the dsRNA duplex via steric hindrance.

### In vivo antitermination and in vitro RNA unwinding activities of putative RBPs

Antitermination activity of proteins can be examined *in vivo* using the RL211 strain of *E. coli* (*46, 57–60*). The chloramphenicol (Cm) acetyltransferase gene (*cat*) is preceded by the Rho-independent *trpL* transcriptional terminator stem-loop that efficiently terminates transcription of the *cat* gene resulting in Cm susceptibility. Expression of a protein with antitermination activity that can unwind or remodel the terminator stem loop results in transcription of the *cat* gene and resistance to Cm (Fig. 5A). We assayed the antitermination activity of our cohort of putative RBPs *in vivo* using the RL211 strain. The RL211 strain was transformed with plasmids harboring the genes of the RBPs under control of the *lac* promoter. Cultures were grown overnight in the absence of Cm and then spotted onto plates with and without Cm plus IPTG (Fig. 5B). All strains grew similarly in the absence of Cm and RL211 did not grow on Cm. Substantial growth on Cm was observed in RL211 strains expressing *stpA*, *bosR*, *cheY2*, *bb0713*, and *flgV* demonstrating their *in vivo* antitermination activity. RL211 strains expressing *ebfC*, *frr*, *bb0796*, and *bpur* grew modestly on Cm plates, while no growth was observed in strains expressing *khpA*, *bb0749* and *grpE*.

**Figure 5.**
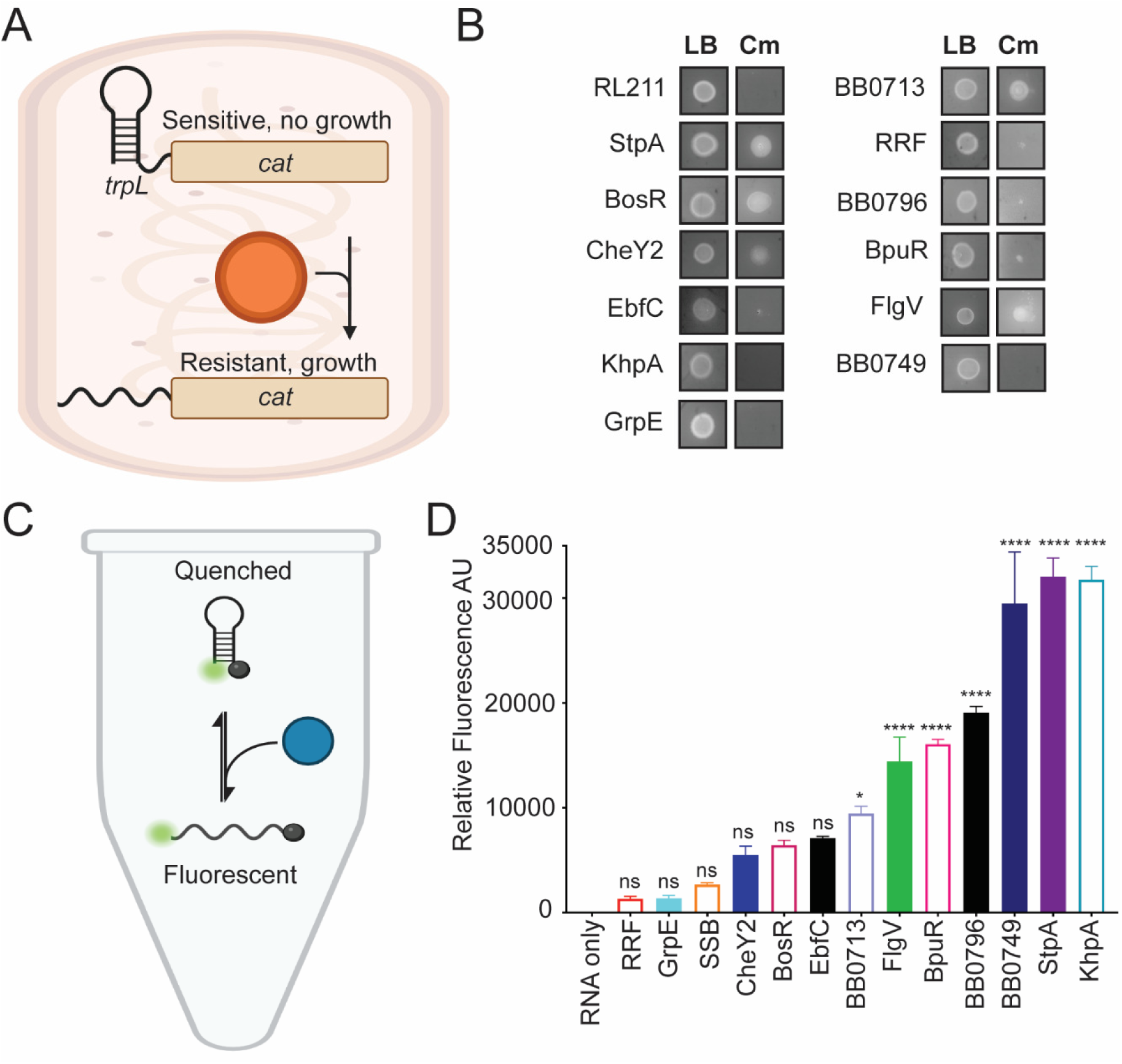
*In vivo* RNA anti-termination and *in vitro* RNA unwinding assays. (A) Illustration of the *in vivo trpL* terminator assay. The *trpL* terminator stem loop is fused to the chloramphenicol (Cm) acetyltransferase (*cat*) gene. In the RL211 strain, the terminator stem loop is formed, transcription is terminated, and the *cat* gene is not expressed resulting in Cm susceptibility and no growth on Cm. However, an RNA chaperone (orange circle) expressed in RL211 that unwinds the terminator stem loop will allow transcription elongation to proceed and the *cat* gene to be expressed, resulting in Cm resistance and growth on Cm (Van Gundy *et al.* 2023). (B) *E. coli* RL211 strains expressing denoted proteins or the positive control StpA were grown in LB overnight to an OD600 of ∼1. The overnight cultures were plated on LB (no antibiotic) or Cm plus 1 mM IPTG, and growth was assessed. (C) Schematic of the *in vitro* unwinding assay. A truncated *trpL* terminator RNA was 5′ and 3′ labeled with 6-FAM and Dabcyl respectively. Protein was added to the *trpL* fluorescent reporter RNA and incubated for 10 min. Florescence was measured to quantify RNA unwinding. (D) A bar graph illustrating the relative fluorescence units of each reaction after subtracting the background levels of unwinding measured in the RNA only reaction. Error bars represent the SEM of three replicates. Relative fluorescent units of unnormalized data were analyzed by a two-way ANOVA with multiple comparisons (p-value ≥0.05, ns, p-value, ≤0.05, p-value ≤0.0001, ****).

The antitermination activity observed in the RL211 strain is likely due to the expressed protein remodeling, unwinding or melting the *trpL* terminator stem loop. We developed a fluorescent reporter to assay the ability of the proteins to remodel or unwind the *trpL* terminator *in vitro*. The *trpL* terminator was 5ʹ and 3ʹ labeled with 6-FAM and Dabcyl, respectively. The labeled *trpL* terminator RNA was incubated with the putative RBPs, and fluorescence was measured to assay RNA unwinding or remodeling (Fig. 5C). BpuR, FlgV, KhpA, BB0713, BB0749, and BB0796 exhibited statistically significant stem-loop remodeling *in vitro* (Fig. 5D). In contrast, BosR, CheY2, EbfC, RRF, SSB, and GrpE did not demonstrate significant RNA remodeling activity compared to RNA alone under the same conditions.

The *in vivo* and *in vitro* data correlate well for BpuR, FlgV, BB0796, and StpA. Both StpA and FlgV exhibited substantial growth on the Cm plates and effectively remodeled the *trpL* terminator *in vitro*. While BpuR and BB0796F showed modest growth on Cm plates and remodeled the *trpL* terminator *in vitro*. In addition, our negative control GrpE did not demonstrate *in vivo* antitermination activity or *in vitro* unwinding activity (Fig. 5). BB0713, BosR and CheY2 exhibited strong *in vivo* antitermination activity as demonstrated by substantial growth on Cm plates. However, the *in vitro* assay revealed that BB0713 only moderately unwound the *trpL* terminator, while BosR and CheY2 showed no statistically significant unwinding of the *trpL* terminator (Fig. 5). BB0713 and BosR both contain two CXXC motifs in their *C*-terminal domains, which often coordinate Zn^2+^ and can be involved oligomerization of proteins. BosR has been reported to bind both DNA and RNA (*40, 61, 62*), and DNA binding requires dimerization (*63*); however, the role of the dimer in RNA binding has not yet been investigated. Since *bosR* is required for borrelial infection (*64, 65*), the data presented here suggests a role for RNA binding by BosR in borrelial pathogenesis. The discrepancy between the activity of BosR in the two assays may be explained by the presence of DTT in the *in vitro* assay buffer, which may influence folding, oligomerization and/or Zn^2+^ binding of BosR, resulting in inhibition of RNA binding and potentially explaining the discrepancies. The RNA annealing and strand displacement assays also utilize the same buffer containing DTT, and the modest activity demonstrated in these assays by BosR and BB0713 may be due to inhibition of oligomerization or Zn^2+^ coordination and proper folding of the proteins. Prior studies demonstrated that a loss of the cysteine residues in the CXXC motifs alters the oligomeric structure of BosR (*63*). In addition, the CXXC domains may serve as redox sensors that affect the binding activity of BosR to both RNA and DNA (*63*). A deeper understanding of the role of tertiary and quaternary structure in these activities awaits further investigation.

Posttranslational modifications (PTMs) of RBPs, including phosphorylation, acetylation, and methylation, regulate their interactions and functions (*4, 5, 66, 67*). In *B. burgdorferi*, CheY2 is phosphorylated by CheA and is postulated to act as a transcriptional or posttranscriptional regulator (*48*). Thus, phosphorylation may act as a molecular switch regulating the RNA chaperone activity and RNA binding of CheY2. CheY2 displayed strong *in vivo* antitermination activity, but did not unwind the *trpL* terminator *in vitro*. We postulate that the phosphorylation state of CheY2 might differ between the purified recombinant protein and the *in vivo* expressed protein in RL211, leading to the observed discrepancy. EbfC and RRF exhibited modest growth on Cm plates, but did not unwind the *trpL* terminator *in vitro*, suggesting that the *in vitro* unwinding assay conditions may be missing important factors found *in vivo*. Notably, the *trpL* terminator we designed for the assay does not include any single-stranded regions other than the stem loop and thus the structure differs from the stem loop *in vivo*. The proteins that performed well in the *in vivo* assay and poorly in the *in vitro* assay may require a toehold or single stranded region before the stem loop in order to bind and unwind the RNA.

RL211 strains expressing BB0749 and KhpA did not grow on Cm plates but unwind the *trpL* terminator robustly *in vitro*. Unfortunately, we cannot determine if the proteins are being properly expressed in the RL211 strain while growing on the plates and the lack of *in vivo* antitermination activity of these proteins may be due to poor protein expression or toxicity due to overexpression of the protein.

### BB0749 binds RNA in B. burgdorferi

Evidence of *in vivo* RNA binding by newly identified RBPs is typically required to classify them as bona fide RBPs. Crosslinking co-immunoprecipitation (CLIP) of the protein bound to RNA is often used to demonstrate *in vivo* RNA binding. However, antibody to the protein or an epitope-tagged version of protein is required for CLIP, therefore high throughput screening by CLIP is not feasible. An *N*-terminal 3×FLAG tagged BB0749 fusion protein was constructed and inserted into the *bb0749* genomic location of the linear chromosome. The *bb0749::3*×*flag* strain was grown to mid-log phase (5 × 10^7^ cells ml^-1^) in one-liter batches and split in two; half of the culture was UV crosslinked while the other half was left untreated as a control. BB0749-RNA complexes were immunoprecipitated with anti-FLAG antibodies and washed several times to remove non-crosslinked RNA from the protein. Western blot analyses demonstrate that BB0749 was enriched and pulled down in the IP in both the crosslinked and non-crosslinked samples (Fig. 6A). However, substantially more RNA was immunoprecipitated with BB0749 in the crosslinked samples compared to the non-crosslinked samples demonstrating that BB0749 binds RNA *in vivo* (Fig. 6B). BB0749 harbors an IDR containing stretches of basic and acidic residues, as well as polar residues. IDRs can directly engage in protein-protein interactions and RNA binding. In eukaryotic cells, IDRs are important for the formation of different types of RNPs, and posttranslational modifications of IDRs modulate their function (*5, 66–68*). IDRs have also been implicated in modulating liquid-liquid phase separation and formation of membrane-less organelles (*69*). Further work is needed to determine the RNA-dependent function of BB0749 and the BB0749 targetome.

**Figure 6.**
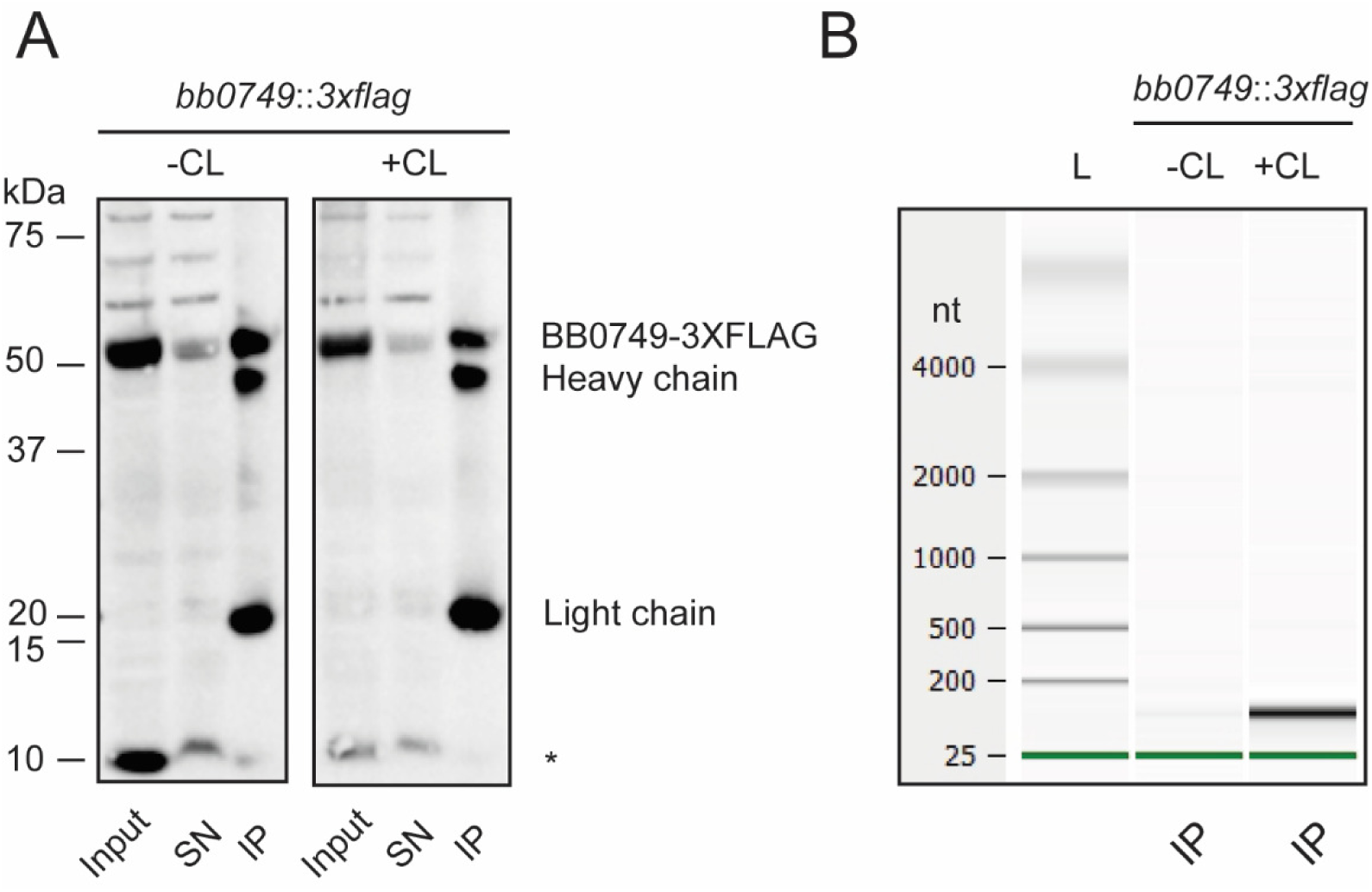
BB0749-3×FLAG binds RNA in *B. burgdorferi*. Crosslinking immunoprecipitation (CLIP) of BB0749-3×FLAG demonstrates RNA binding *in vivo*. (A) *N*-terminally tagged BB0749 cultures were grown and split in half. One half was UV crosslinked (+CL), while the other was not (-CL). The cell lysates from the two samples were immunoprecipitated with anti-FLAG antibody. (A) Cell lysates (input), supernatants after the immunoprecipitation (SN) and the immunoprecipitations (IP) were separated on a denaturing SDS polyacrylamide gel and transferred to a PVDF membrane and immunoblotted with anti-FLAG antibody. BB0749 (∼50 kDa) is detected in all samples but is depleted in the SN and enriched in the IP. Preliminary mass spectrometry data suggest the smaller band detected and denoted with an * may be a small isoform or proteolytic fragment of BB0749 (∼10 kDa). The heavy and light chain of the anti-FLAG antibody beads are also detected (∼22 kDa and ∼50 kDa). (B) Bioanalyzer gel image of non-crosslinked (-CL) and crosslinked (+CL) IP RNA samples. Green lines represent bioanalyzer lower marker. All experiments were performed in triplicate.

### FlgV has RNA chaperone activity

FlgV (BB0268), previously termed HfqBb (*45*), anneals and strand displaces RNA in both the gel-based and in-solution assays. In addition, FlgV had antitermination activity *in vivo* and RNA unwinding *in vitro* (Table S5). Fifteen years ago, we identified BB0268 as an RBP that partially complemented *rpoS* posttranscriptional regulation in an *hfq* deletion mutant in *E. coli* and we showed *E. coli hfq* partially complemented the *bb0268* deletion mutant in *B. burgdorferi*. We termed BB0268 HfqBb because of the reciprocal complementation of the deletion phenotypes and demonstrated BB0268 bound RNA non-specifically *in vitro* (45). Recently, Zamba-Campero *et al.* reported that BB0268 monomer is a flagellar protein and the Alphafold3 predicted structure of BB0268 is not similar to the structure of Hfq (*44*). BB0268 was renamed FlgV and we agree that it was misannotated as HfqBb. However, Zamba-Campero *et al.* also concluded that FlgV was not an RBP because RNA did not Co-IP with FlgV after UV crosslinking (*44*). Successful UV crosslinking relies on spatial arrangement and proximity of the RNA and protein, as well as the nucleotide and amino acid composition at the RNA protein interface (*70*). Protein interactions with the phosphate backbone of RNA are suboptimal and are likely undetected by UV crosslinking. In addition, UV crosslinking efficiency is poor, approximately 1-5%; abundant and/or stable RNA-protein complexes are more likely to be captured than rare and/or transient complexes using UV crosslinking immunoprecipitation (CLIP) (*70*). Despite these caveats, Zamba-Campero *et al.* conclude that FlgV is not an RBP because levels of bulk RNA recovered after CLIP with FlgV do not match the levels of RNA recovered from two known RBPs, *E. coli* Hfq and *B. burgdorferi* KhpB (*44*). A direct comparison of the ability of these proteins to co-immunoprecipitate RNA is not feasible and, in fact, misleading, due to inherent structural differences between the proteins and the distinct mechanisms of RNA binding. Generally, instead of comparing RNA from Co-IP’s of different proteins to determine RNA binding, a non-crosslinked sample is compared to the crosslinked sample to determine if the RNA observed in the co-IP is specific, as demonstrated for BB0749 (Fig. 6). The study also included CLIP of endogenous FlgV with a polyclonal antibody produced against FlgV (*44*). Again, a non-crosslinked control was not included, but a co-IP was performed on a *bb0268* null mutant. The profiles of the RNA immunoprecipitated from the mutant is different than the wild-type strain, but the RNAs were not sequenced to determine if any specific RNA was pulled down with FlgV. In both cases, a transient interaction or an interaction with low abundance RNA would not be detected by comparing bulk RNA amounts and patterns.

FlgV forms a monomer and a dimer *in vitro* and *in vivo* (*44, 45*). While the monomer clearly plays a role in flagellar structure, the dimer was not observed in association with the flagella apparatus and its function was not discussed (*44*). The data we presented previously and in this study demonstrate that FlgV has RNA chaperone activity and non-specific RNA binding *in vitro* (*45*). We hypothesize that FlgV dimer (and possibly monomer) moonlight as an RNA-binding protein. Moonlighting, enigma or unconventional RBPs are involved in biological processes with no apparent relation to RNA biology: thousands of unconventional RBPs have been identified (*5, 6, 8, 10*). Several dozen moonlighting RBPs have been validated by orthogonal approaches including co-IPs in eukaryotic cells. Notably, flagellar and flagellar associated proteins (FlhA, FliN, FliH, FlgN, FliL, FliS, and YgcR) were identified as RNA-binding proteins in *E. coli* using total RNA-associated protein purification (TRAPP) (*8*). TRAPP is based on UV crosslinking of RNA and RBPs in cells followed by purification of bulk RNA on silica beads under denaturing conditions. Proteins that are not crosslinked to RNA are washed away and proteins covalently crosslinked to RNA remain bound and purified with the RNA. *In vivo* identification of flagellar and flagellar-associated proteins as RBPs using RNA-centric approaches in other bacteria suggest flagellar proteins can moonlight as RBPs (*8*).

### Newly identified proteins with RNA chaperone activity

The primary objective of this study was to identify previously uncharacterized RNA binding proteins (RBPs) with RNA chaperone activity in *B. burgdorferi*. Among the proteins tested, uncharacterized proteins BB0713, BB0749 and BB0796 demonstrated robust RNA chaperone activity, yielding positive results in five of the six functional assays employed (Table S5). BB0713 and BB0749 both have domains that have been reported to bind RNA (Zn knuckle and IDR, respectively), while BB0796 does not. The Alphafold3 predicted structure of BB0796 has homology to the Skp (OmpH) family of proteins, which includes a coiled-coil domain. Skp has been reported as a molecular chaperone that interacts with unfolded proteins as they emerge in the periplasm in *E. coli*. In yeast, heat shock 70 kDa protein 1 (Hsp70-1) and Hsp90 domains, which also interact with unfolded and misfolded proteins, were reported to bind RNA *in vivo*. In addition, CheY2 also demonstrated RNA chaperone activity in five of six functional assays (Table S5). Although CheY2 is annotated as a chemotaxis response regulator due to its sequence and structural homology to other chemotaxis response regulators, it is not necessary for chemotaxis or motility in *B. burgdorferi*. However, CheY2 is important for vertebrate infection and was postulated to be a transcriptional or posttranscriptional regulator of virulence gene expression (*48*). Our data suggest CheY2 may regulate gene expression via its RNA chaperone activity. Notably, CheY and other chemotaxis associated proteins (CheW, CheZ, CheB, and Trg) were identified as RNA binding proteins via TRAPP in *E. coli* (*8*). Chemotaxis proteins moonlighting as RNA binding proteins may be a conserved function of these proteins in bacteria.

### Concluding remarks

We identified approximately 10% of the annotated *B. burgdorferi* proteome in our mass spectrometry analysis of the first ten fractions of the gradient-separated cell lysate. The coverage of the proteome was less than what has been reported in Grad-seq for other bacteria (*11–17*). The lower coverage may be due to limited input material, technical differences in preparing fractions, or not partitioning of the fractions before mass spectrometry. In addition, gradient profiling is not efficient at capturing low abundant and transient RNA-protein interactions. The RNPs are not crosslinked, and transient or unstable complexes can disassociate after cell lysis and during gradient fractionation. There are likely RBPs that were not identified in our assay due to these caveats. Global RNA-centric approaches, such as 2C or TRAPP, or protein-centric approaches are needed to further identify RBPs in *B. burgdorferi*. Notably, we found most of the known RBPs in gradient profiling and identified several novel RBPs; in addition, we characterized the *in vitro* RNA chaperone activity of many of these RBPs. Further investigation is necessary to elucidate the cellular functions and RNA interactomes of both previously characterized and newly identified RBPs. RBPs are integral to numerous cellular processes and serve as critical regulators of gene expression, with significant implications for pathogenesis and virulence. The identification and functional characterization of novel RBPs in *B. burgdorferi* will enhance our understanding of the molecular mechanisms that facilitate spirochete transmission, host adaptation, and disease pathogenesis.

## Materials and methods

### Bacteria and strains

*B. burgdorferi* low-passage virulent B31 strain (clone B31-5A4) was cultivated in Barbour-Stoenner-Kelly II (BSK-II) complete medium at 37°C. Unless otherwise stated, cultures were passaged to 1 × 10^5^ cells ml^-1^ and grown to 1-3 × 10^7^ cells ml^-1^. Cell density was determined using a Petroff Hausser counting chamber as previously described (*71, 72*). *E. coli* RL211 strain constructed by Landick *et al.* (*57*) was cultivated in lysogeny broth (LB) at 37°C and kanamycin (50 ug/ml) when appropriate.

### Gradient fractionation

The protocol was adapted from Smirnov *et al.*, 2016 (*11*). *B. burgdorferi* B31-5A4 cultures were centrifuged at 5,000 × g for 30 min at 4°C and the cell pellets were snap frozen in liquid nitrogen (LN2). In total 5 × 10^8^ to 1.5 × 10^9^ cells were lysed in 100 µl of lysis buffer (20 mM Tris-HCl, pH 7.5, 150 mM KCl, 1mM MgCl2, 1mM DTT, 1mM PMSF, 0.2% Triton X-100, 20 units/ml DNase I (Thermo-Fisher Scientific™), 200 units/ml SUPERase-IN (Life Technologies)) with a stainless-steel ball cryogrinding (Retsch) in the Retsch MM400 at 30 Hz for 10 min, refreezing the sample in LN2 every 3 minutes. The lysate was cleared via centrifugation at 18,928 × *g* for 5 min at 4°C. Cleared lysate was applied to a 10-40% (wt/vol) glycerol gradient and ultra-centrifuged in a linear Beckman SW40Ti tube at 62,909 × *g* for 17 h at 4°C. The gradient was separated into 20 equal fractions of 600 µL and approximately 90 µL of each fraction was removed for protein analyses. The remaining sample was used for RNA isolation.

### Protein gel electrophoresis

Gradient fractions (10 µL of each fraction in Laemmli buffer) were resolved on 12% SDS-PAGE at 150V for 1 h at room temperature followed by Coomassie Brilliant Blue staining, overnight. The gel was destained with distilled water for 4-10 h, changing the water approximately every thirty minutes.

### RNA isolation and gel electrophoresis of gradient fractions

RNA was isolated from approximately 500 µL of each fraction. Each fraction was treated with 1% SDS and one volume of saturated phenol (pH 4.3) and incubated at 55°C for 5 min, centrifuged 18,928 × *g* for 10 min at 4°C and the aqueous layer moved to a new tube. RNA was precipitated with 0.3M NaOAc and three volumes of ice-cold absolute ethanol at -20°C overnight. RNA was pelleted for 45 min at 18,928 × *g* at 4°C and washed with 75% ice-cold ethanol. RNA was centrifuged at 18,928 × *g* at 4°C for 20 min, the supernatant was removed, and the pellet was air-dried and resuspended in water. RNA was resolved on a denaturing 6% polyacrylamide gel with 7M Urea in 1× TBE buffer. The gel was post stained with ethidium bromide.

### Mass-spectrometry sample preparation and analysis

Glycerol gradient fractions were diluted with lysis buffer (4% (w/v) SDS, 0.1M Tris pH 7.4) and reduced with 5mM TCEP at 90°C for 10 minutes. Samples were cleared by centrifugation and transferred to a 30kDa spin filter (Amicon). Samples were washed twice with UA buffer (8M Urea, 0.1M Tris pH 7.4) then alkylated with 50mM iodoacetamide at room temperature for 20 minutes in the dark. Samples were washed again three times with UA and then three times with 0.1M ammonium bicarbonate (ABC), 0.02% (w/v) deoxycholic acid (DA) then suspended in 150 µL 0.1M ABC, 0.02% DA with trypsin overnight at room temperature. Tryptic peptides were eluted and DA was removed by phase transformation using ethyl acetate. Tryptic peptides were then desalted using Pierce C18 spin columns according to the manufacturer’s instructions and dried with a vacuum centrifuge. Tryptic peptides were loaded directly onto a Waters nanoACQUITY UPLC BEH C18 column (130 Å, 1.7 µm, 75 µm x 250 mm) and separated with a nanoAcquity UPLC (Waters) at 0.3 µl/min using gradients of 3 to 8% acetonitrile in 5 minutes then 8 to 35% acetonitrile in 118 minutes. Detection of peptides was performed with a LTQ Orbitrap Velos mass spectrometer (Thermos Scientific), scanning from 300 to 1800 m/z at 60,000 resolution with an AGC target of 1E6 and selecting the 10 most intense ions for MS/MS. Maximal ion injection times were 500 ms for FT (one microscan) and 250 ms for LTQ with an AGC target of 1E4. The normalized collision energy was 35% with activation Q 0.25 for 10 ms. Raw data files were searched against the Uniprot *Borreliella burgdorferi* proteome database, using the MaxQuant/Andromeda search engine (version 2.6.3.0). Trypsin cleavage was searched allowing for up to two missed cleavages, with cysteine carbamidomethylation as a fixed modification, and protein *N*-terminus acetylation and methionine oxidation as variable modifications. Mass tolerances were set to 20 ppm (first search) and 4.5 ppm (main search) for precursor ions, and 0.5 Da for ITMS MS/MS ions. All peptides and proteins were thresholded at 1% FDR. The mass spectrometry proteomics data have been deposited to the ProteomeXchange Consortium via the Proteomics Identifications Database partner repository with the dataset identifier PXD068462 and 10.6019/PXD068462

### Northern blotting

RNA was separated under denaturing conditions on pre-poured 6% Novex™ TBE-Urea (7M) gels (Invitrogen™) in 1× TBE. RNA was denatured at 65°C for 15 min in 2× RNA load dye (Thermo-Fisher Scientific™) prior to loading on the gel. RNA was transferred to BrightStar™-Plus positively charged nylon membrane by electroblotting at 10V for 1 h in 0.5× TBE. Membranes were UV crosslinked and probed with biotinylated RNA probes. A biotinylated 6S probe was generated from *in vitro* transcription of PCR-generated DNA template using the MAXIscript™ T7 Transcription Kit (Invitrogen™) with biotinylated-16-UTP (Roche) according to the manufacturer’s protocol. The T7 bacteriophage promoter was added to the 5ʹ end of the reverse primer that generated the 6S DNA template (Table S3). Probe length and integrity were examined by resolving the in vitro transcription reaction on a pre-poured 6% Novex™ TBE-Urea (7M) gel (Invitrogen™) in 1× TBE and post staining with ethidium bromide. The remainder of the reaction was ethanol precipitated and used for Northern blotting. Membranes were pre-hybridized in 4 ml of ULTRhyb™ Ultra-sensitive Hybridization Buffer (Invitrogen™) at 68°C for 30 min in 50 ml conical tubes in a hybridization oven with rotation. Subsequently, 500 µL of the pre-hybridization solution was transferred to 1.5-ml microfuge tube and the probe was added to the pre-hybridization solution to a final concentration of approximately 0.1 nM. The pre-hybridization buffer with probe was incubated at 95°C for 5 min, immediately returned to the prehybridizing membrane and incubated overnight at 68°C with rotation. Membranes were washed for 5 min in 2× SSC and 0.1% SDS at room temperature with rotation and twice for 15 min in 0.1× SSC, and 0.1% SDS at 68°C with rotation. Membranes were developed using the Chemiluminescent Nucleic Acid Detection Module Kit (Thermo-Fisher Scientific™) according to the manufacturer’s protocol and visualized on an Azure Sapphire Biomolecular Imager.

### Immunoblotting

Each fraction was diluted in 2× Laemmli buffer (Sigma-Aldrich), heated to 95°C for 5 min and resolved on precast Criterion Any kD™ TGX Stain-Free™ Protein Gels (BioRad). Gels were semi-dry blotted onto prepackaged Iblot™ 2 Transfer Stacks, PVDF membrane (Invitrogen™) using a custom Iblot2 program (20 V for 3 min, 23 V for 1 min, 25 V for 1 min). Membranes were blocked for 1 h in blocking buffer [1.5 g skim milk 1× TBST (1.5 M NaCl, 10 mM Tris–HCl, pH 7.5, 0.05% Tween 20)] (Thermo-Fisher Scientific™). Protein-specific antisera were used to detect RNA polymerase β subunit, BosR and BB0749 proteins in the gradient fractions. The mouse anti-RNA polymerase β subunit antibody was purchased (Biolegend), as was the anti-BosR antibody (General Biosciences), and rat anti-BB0749 was generated in this study. Primary antisera were diluted 1:20,000 in 1× TBST and incubated for 1 h at room temperature. Membranes were washed 3 × 5 min in 1× TBST. Secondary antibodies, goat anti-mouse HRP or goat anti-rat HRP (KPL) were diluted 1:20,000 in 1× TBST and incubated for 1 h at room temperature. Membranes were developed using chemiluminescence (ECL select, Amersham) and imaged on an Azure Sapphire biomolecular imager using the chemiluminescence setting with High Dynamic Range (HDR), auto exposure, and single capture.

### BB0749 recombinant protein purification and antisera production

Recombinant protein was generated as previously described in Van Gundy *et al.*(*46*). All animal studies were conducted using the Guide for the Care and Use of Laboratory Animals (Institute of Animal Research, National Research Council) with protocols approved by the Virginia Commonwealth University IACUC. Eight-week-old female Sprague-Dawley rats (Charles River) were immunized on a three-dose schedule, with each dose two weeks apart as previously described (*73*). Rats were primed by IP vaccination with 50 µg antigen, BB0749 recombinant protein, in Freund’s complete adjuvant (Sigma-Aldrich). Two boosters of 25 µg antigen, BB0749 recombinant protein, each in Freund’s incomplete adjuvant were also administered IP (Sigma-Aldrich). One week following the final boost, rats were euthanized by CO2 asphyxiation and terminal bleeds conducted by cardiac puncture. Serum was harvested by centrifugation in VACUETTE Z serum sep-clot activator tubes (Greiner Bio-One) according to manufacturer instructions.

### Recombinant protein expression and FPLC purification

Plasmids for recombinant protein production were constructed using GenScript Plasmid Synthesis service. Open reading frames were codon optimized for *E. coli*. The codon optimized open reading frames and final plasmid sequences used can be found in Table S3. Plasmids were purified using PureYield™ Midiprep plasmid system (Promega) per manufacturer’s protocol. Plasmids were transformed into *E. coli* BL21(DE3) cells (Lucigen™) and transformants were screened by PCR using pETite T7 Forward Primer and pETite Reverse Primer (50 µM). Positive transformants were grown in LB containing kanamycin (50 μg/ml) overnight. The overnight culture was inoculated 1:100 into LB supplemented with kanamycin (50 μg/ml) and grown to mid-logarithmic phase (OD600 = 0.4-0.6). Protein expression was induced by the addition of isopropyl-D-thiogalactopyranoside (IPTG) to a final concentration of 1 mM and incubated with shaking for 16-18 hours at 20°C. Cultures were centrifuged at 5000 × g for 10 min. The cell pellets were resuspended in buffer A (20 mM NaH2PO4, 500 mM NaCl, 20 mM imidazole, pH 7.4) containing 1mM MgCl2, Universal Nuclease for Cell Lysis (10U/ml, Pierce™ Thermo-Fisher Scientific™), and 0.2 mg/ml lysozyme and incubated for 30 min and vortexed every 10 min. The lysate was centrifuged at 12,000 × *g* for 30 min at 4°C to separate the soluble and insoluble fractions. The soluble fractions were passed through a 5 μM filter (Millipore) and proteins were purified using nickel affinity chromatography by a Fast Protein Liquid Chromatography platform (ÄKTA Pure 25 M; Cytiva) with a 1-ml HisTrap FF column (Cytiva) at a flow rate of 1 ml/min. Columns were equilibrated using five column volumes of buffer A. Samples were loaded mechanically, followed by washing with 20 column volumes of buffer A. Proteins were eluted in 1 ml increments with five column volumes of elution buffer (20 mM NaH2PO4, 500 mM NaCl, 500 mM imidazole). Fractions were combined and dialyzed in G3 Slide-a-lyzer™ cassettes (Thermo-Fisher Scientific™, 3.5 MWCO, 3 ml cassette) in 1× PBS (phosphate-buffered saline, pH 7.4). Dilute proteins were concentrated with Pierce Concentrators, PES, 3K MWCO, 0.5 ml (Thermo-Fisher Scientific™). Proteins were quantified by bicinchoninic acid (BCA) assay (Thermo-Fisher Scientific™), resolved on precast Criterion Any kD™ TGX Stain-Free™ Protein Gels (BioRad) and visualized by silver staining to assess protein purity (Fig. S7).

### RNAs

RNAs were ordered from Dharmacon or IDT, dissolved in water and stored at -20°C. Sequences are listed in Table S3.

### In-solution RNA annealing and strand displacement assay

The combined RNA annealing and strand displacement assay was adapted from the Schroeder laboratory but utilizes fluorophores and quenchers instead of fluorescence resonance energy transfer (FRET). Two previously published 21-nucleotide complementary RNAs were used with 5ʹ and 3ʹ modifications (*46, 54*). M1 RNA included a 5ʹ 6-FAM fluorophore (M1_5FAM6), J1 RNA included a 3ʹ Dabcyl quencher (J1_3Dab), and the unlabeled competitor uJ1 contained no modifications (Table S3). Combined RNA annealing and strand displacement assays were carried out in 1× FRET buffer (50 mM Tris-HCl pH 7.5, 1 mM MnCl2, 10 mM NaCl and 1 mM DTT) at 37°C unless otherwise stated. Notably the 1× FRET buffer contains manganese instead of magnesium. Manganese was reported to be important for *B. burgdorferi* survival and proper function of RNAP *in vitro* (*74–76*). In-gel annealing and strand displacement assays in 1× FRET buffer with 1 mM MnCl2 or 1 mM MgCl2 were performed, and the rate of strand displacement was faster with Mn instead of Mg, while annealing kinetics were similar between both salts (Fig. S8). Final concentrations of M1_5FAM6 and J1_3Dab RNA in each reaction were 25 nM and the unlabeled competitor M1 RNA (uM1) was added to a final concentration of 250 nM. Final concentrations of protein in the reaction were 1 µM. Proteins were stored in 1× PBS, therefore an equal volume of 1× PBS was added to all reactions, including the negative control RNA only reaction. 100 µl combined RNA annealing and strand displacement reactions were carried out in Nunc™ Microwell™ black flat-bottom 96-well plates (Thermo-Fisher Scientific™) and read on a Tecan Spark. The annealing reaction was started by injecting J1_3Dab into wells containing M1_5FAM6 in 1× FRET buffer with or without protein. The reaction was shaken for 3 s and the samples were excited at 485 nm (20 bandwidth), and emission intensity was measured at 535 nm (25 nm bandwidth) every 10 s for 180 s. The stand displacement assay was then initiated by injecting tenfold molar excess of unlabeled M1 (uM1). The reaction was shaken for 3 s and readings were taken every 10 s for 180 s. At least three reactions were performed for each protein. The annealing reaction final time point was constrained to zero by subtracting the fluorescent units at the final time point from itself. Accordingly, all timepoints’ fluorescent units were normalized by subtracting the zero time point fluorescent units from them. In addition, the initiation of the annealing reaction was constrained to one by dividing the fluorescent units of the timepoint before injection of J1 by itself, and all timepoints’ fluorescence units were normalized by dividing them by the initiation time point fluorescent units. These normalizations constrain the data from 0 to 1 (Fig. S3), which was used in the GraphPad Prism statistical analyses. Annealing rates were measured using one-phase decay and strand displacement rates were measured using one phase association.

### Gel-based RNA annealing and strand displacement assays

The RNA gel annealing and strand displacement assay protocols were adapted from the Schroeder laboratory and performed as previously described (*46*).

### Anti-termination assay in *E. coli*

Transcription anti-termination assays were performed as described (*46*). *E. coli* RL211 strain containing the *cat* gene preceded by the *trpL* terminator stem-loop were transformed with plasmids constructed for purifying recombinant proteins (described above and in Table S3). Cells were grown overnight in LB and when appropriate kanamycin (50 µg/ml) was added. Overnight cultures were spotted (5 µL) on LB and LB with chloramphenicol (34 µg/ml) plates and grown at 37°C. Cell growth was inspected daily. Plates were imaged using the Azure Sapphire biomolecular imager using the white light channel. Experiments were repeated in triplicate.

### *In vitro* RNA unwinding assay

The truncated Rho-independent *trpL* terminator fluorescent reporter RNA contained a 5ʹ 6-FAM fluorophore and a 3ʹ Dabcyl quencher (*trpL* term). A 100 nM working stock of *trpL* term in 1× FRET buffer was heated to 65°C for 10 min, slow cooled to 37°C and stored on ice. Reactions contained 50 nM *trpL* term and 1 µM protein (final concentrations). Proteins were stored in 1× PBS, therefore an equal volume of 1× PBS was added to all reactions, including the negative control RNA only reaction. 100 µl unwinding reactions were carried out in Nunc™ Microwell™ black flat-bottom 96 well plates (Thermo-Fisher Scientific™) buffer at 24°C using the Tecan Spark. The unwinding reaction was started by injecting the *trpL* term RNA into wells containing 1× FRET buffer (50 mM Tris-HCl, pH 7.5, 1 mM MnCl2, 10 mM NaCl and 1 mM DTT) with or without protein and incubated for 10 min. The samples were excited at 485 nm (20 bandwidth), and emission intensity was measured at 535 nm (25 nm bandwidth). At least three reactions were performed for each protein. Data was background normalized by removing the background fluorescence signal (RNA without protein) from all samples. Dunnett’s Multiple Comparison Two Way ANOVA was used to compare each protein against RNA only (ns, p<.05,*, p<.005,**, p<.0001,****).

### *Bb0749::3×flag* strain construction

Low-passage virulent *B. burgdorferi* strain B31-5A4 genomic DNA was used as template for amplifying the *bb0749* upstream region and *bb0749* ORF using Phusion™ High Fidelity DNA polymerase and region-specific primers (Table S3). The PCR products were cloned into pGEM®-T Easy (Promega) per manufacturer’s instructions and transformed into DH5α *E. coli* cells (Thermo-Fisher Scientific™). A gBlock™ was synthesized by IDT that contained the streptomycin/spectinomycin resistance cassette (*flgBp-aadA)*, the *bb0749* predicted promoter and 5′ UTR, and the *B. burgdorferi* optimized *3×flag* sequence (Table S3). The gBlock DNA was used as template to amplify the sequence using Phusion™ High Fidelity DNA polymerase and gBlock specific primers (Table S3). The PCR product was cloned into pGEM®-T Easy (Promega) per manufacturer’s instructions. Positive transformants were validated by sequencing. The *bb0749* ORF region 5′ end was engineered with an XmaI and HindIII site and the 3′ end was engineered with an AscI site. The gBlock 5′ end was engineered with an XmaI site and the 3′ end was engineered with a HindIII site. The NdeI site found between the *flgB* promoter sequence and the *aadA* ORF of the streptomycin/spectinomycin antibiotic resistance cassette was mutated and an NdeI site was added between the 3′ end of *bb0749* 5′ UTR and the *3×flag* sequence. The gBlock was inserted into the *bb0749* ORF construct with XmaI and HindIII yielding the gBlock::ORF construct. Positive transformants were sequence validated. The *bb0749* upstream region 3′ end was engineered with an XmaI and AscI site, which was used to insert the gBlock::*bb0749* ORF into the upstream region, yielding *bb0749*up::gBlock::ORF. Positive transformants were validated by sequencing. The final plasmid: pGEM®-T Easy *bb0749* up:gBlock:ORF was linearized with AhdI for *B. burgdorferi* transformation, as described previously (*71*). Positive transformants were screened with primers flanking the insertion site on the chromosome (Table S3). The *bb0749::3×flag* strain was validated by sequencing and verified by Western blot for expression of BB0749-3×FLAG.

### Crosslinking immunoprecipitation of *bb0749::3×flag*

*B. burgdorferi bb0749::3×flag* strain was cultivated in BSK-II media at 37°C. Cells were grown to 5 × 10^7^ cells ml^-1^, split into two 500-ml cultures and collected by centrifugation. One pellet was treated as the non-crosslinked control and was washed in 2 ml of dPBS, spun at 18,000 × *g* for 2 min, the supernatant was discarded, and the cell pellets were placed at -80^°^C. The crosslinked sample pellet was resuspended in 100 ml of dPBS and 20 ml of the resuspended cells were crosslinked at a time. 20 ml of resuspended cells were placed into a glass baking dish and UV crosslinked twice with 80,000 μJ/cm^2^ in a Spectrolinker^™^ XL-1000 UV Crosslinker (Spectroline^™^). The crosslinked samples were combined and centrifuged at 5000 × *g* for 10 min, the supernatant was discarded, the cell pellet was resuspended in 2 ml of dPBS and centrifuged at 18,000 × *g* for 2 min, the supernatant was discarded. The crosslinked pellet was stored at -80^°^C. Non-crosslinked and crosslinked pellets were thawed on ice and resuspended in 1 ml of ice-cold lysis buffer (50 mM Tris-HCl, pH 7.5, 5 mM MgCl2, 250 mM NaCl, cOmplete protease inhibitor mini tablet, 10 U/µl SupeRNasin.). Cells were placed in 5-ml steel cryo-vials (Retsch), submerged in LN2 and subsequently lysed at 30Hz for 10 min, submerging samples in LN2 every 3 min, in the Retsch MM400. The cell lysate was centrifuged at 18,000 × *g* for 30 min at 4C° and the cleared lysate transferred to a new tube. Input samples were taken for western blot analysis. Cleared lysate was added to 15 µl of anti-FLAG M2 magnetic beads (Sigma-Aldrich) that were equilibrated with lysis buffer. Samples rotated overnight at 4°C and placed on a magnetic stand. The supernatant from each sample was collected for western blot analysis. Samples were washed three times with 500 µL TBS buffer (50 mM Tris-HCl, pH 7.5, 150 mM NaCl). The beads were resuspended in 220 µl and 20 µl were collected for western blot analyses. The sample was placed on the magnetic stand and the supernatant was discarded. The beads were resuspended in Zymo Research Quick-RNA Miniprep Lysis Buffer (Cat #: R1060-1-50) and incubated at RT for 10 min. The samples were placed on the magnetic stand and the supernatant was collected. The RNA was purified using the Zymo Research Quick-RNA^™^ MiniPrep Plus (Zymo Research #R1057) per manufacturer’s instructions. The protocol for the RNA miniprep kit was repeated twice and ethanol was added before the Spin-Away™ Filter to increase the binding of DNA and large RNAs. Isolated RNA samples were run on a Bioanalyzer 2100 Pico chip and protein was visualized via western blot.

## Ethical approval

This article does not contain any studies with human participants. All animal studies were conducted using the Guide for the Care and Use of Laboratory Animals (Institute of Animal Research, National Research Council) with protocols approved by the Virginia Commonwealth University IACUC.

## Author contributions

MCL, TV, CS, JTS, JH, DSS and RM conceived the study. ML, TV, CE, DD, GC, NC, KF performed experiments. TV, MCL, GC and CE wrote the manuscript. All authors read and approved the final manuscript.

## Acknowledgements

We thank Bob Gilmore, Kevin Brandt, Shelby Lennon, Joseph Cardiello, and Marisa Foster for thoughtful and critical reading of the manuscript. We thank Jeremy Bono and James Kovaks for useful discussions. We thank Robert Landick for the RL211 strain. The findings and conclusions in this report are those of the authors and do not represent the official position of the Centers for Disease Control and Prevention.

## References

1. P. Babitzke, Y. J. Lai, A. J. Renda, T. Romeo, Posttranscription Initiation Control of Gene Expression Mediated by Bacterial RNA-Binding Proteins. Annu Rev Microbiol 73, 43–67 (2019).

2. E. Ng Kwan Lim, C. Sasseville, M. C. Carrier, E. Massé, Keeping Up with RNA-Based Regulation in Bacteria: New Roles for RNA Binding Proteins. Trends Genet 37, 86–97 (2021).

3. E. Holmqvist, J. Vogel, RNA-binding proteins in bacteria. Nat Rev Microbiol 16, 601–615 (2018).

4. M. Corley, M. C. Burns, G. W. Yeo, How RNA-Binding Proteins Interact with RNA: Molecules and Mechanisms. Mol Cell 78, 9–29 (2020).

5. M. W. Hentze, A. Castello, T. Schwarzl, T. Preiss, A brave new world of RNA-binding proteins. Nat Rev Mol Cell Biol 19, 327–341 (2018).

6. A. Castello et al., Insights into RNA biology from an atlas of mammalian mRNA-binding proteins. Cell 149, 1393–1406 (2012).

7. J. M. Smith, J. J. Sandow, A. I. Webb, The search for RNA-binding proteins: a technical and interdisciplinary challenge. Biochem Soc Trans 49, 393–403 (2021).

8. V. Shchepachev et al., Defining the RNA interactome by total RNA-associated protein purification. Mol Syst Biol 15, e8689 (2019).

9. C. Asencio, A. Chatterjee, M. W. Hentze, Silica-based solid-phase extraction of cross-linked nucleic acid-bound proteins. Life Sci Alliance 1, e201800088 (2018).

10. L. C. Chu et al., The RNA-bound proteome of MRSA reveals post-transcriptional roles for helix-turn-helix DNA-binding and Rossmann-fold proteins. Nat Commun 13, 2883 (2022).

11. A. Smirnov et al., Grad-seq guides the discovery of ProQ as a major small RNA-binding protein. Proc Natl Acad Sci U S A 113, 11591–11596 (2016).

12. J. Hor et al., Grad-seq in a Gram-positive bacterium reveals exonucleolytic sRNA activation in competence control. EMBO J 39, e103852 (2020).

13. J. Hor et al., Grad-seq shines light on unrecognized RNA and protein complexes in the model bacterium *Escherichia coli*. Nucleic Acids Res 48, 9301–9319 (2020).

14. M. Gerovac et al., A Grad-seq View of RNA and Protein Complexes in *Pseudomonas aeruginosa* under Standard and Bacteriophage Predation Conditions. mBio 12, (2021).

15. M. Gerovac et al., Global discovery of bacterial RNA-binding proteins by RNase-sensitive gradient profiles reports a new FinO domain protein. RNA 26, 1448–1463 (2020).

16. K. Chihara, M. Gerovac, J. Hör, J. Vogel, Global profiling of the RNA and protein complexes of *Escherichia coli* by size exclusion chromatography followed by RNA sequencing and mass spectrometry (SEC-seq). RNA 29, 123–139 (2022).

17. M. Gerovac, J. Vogel, A. Smirnov, The World of Stable Ribonucleoproteins and Its Mapping With Grad-Seq and Related Approaches. Front Mol Biosci 8, 661448 (2021).

18. K. Katsuya-Gaviria, G. Paris, T. Dendooven, K. J. Bandyra, Bacterial RNA chaperones and chaperone-like riboregulators: behind the scenes of RNA-mediated regulation of cellular metabolism. RNA Biol 19, 419–436 (2022).

19. S. A. Woodson, S. Panja, A. Santiago-Frangos, Proteins that chaperone RNA regulation. Microbiology Spectrum 6, RWR-0026-2018 (2018).

20. S. A. Woodson, Taming free energy landscapes with RNA chaperones. RNA Biol 7, 677–686 (2010).

21. L. Rajkowitsch, R. Schroeder, Dissecting RNA chaperone activity. RNA 13, 2053–2060 (2007).

22. L. Rajkowitsch et al., RNA chaperones, RNA annealers and RNA helicases. RNA Biol 4, 118–130 (2007).

23. M. Doetsch, R. Schroeder, B. Fürtig, Transient RNA-protein interactions in RNA folding. FEBS J 278, 1634–1642 (2011).

24. O. Duss, G. A. Stepanyuk, J. D. Puglisi, J. R. Williamson, Transient Protein-RNA Interactions Guide Nascent Ribosomal RNA Folding. Cell 179, 1357–1369.e1316 (2019).

25. K. J. Kugeler, A. M. Schwartz, M. J. Delorey, P. S. Mead, A. F. Hinckley, Estimating the Frequency of Lyme Disease Diagnoses, United States, 2010-2018. Emerg Infect Dis 27, 616–619 (2021).

26. D. S. Samuels et al., Gene regulation and transcriptomics. Current Issues in Molecular Biology 42, 223–266 (2021).

27. A. L. Sapiro et al., Longitudinal map of transcriptome changes in the Lyme pathogen. Elife 12, (2023).

28. A. A. Grassmann et al., BosR and PlzA reciprocally regulate RpoS function to sustain *Borrelia burgdorferi* in ticks and mammals. J Clin Invest 133, (2023).

29. M. Lybecker, I. Bilusic, R. Raghavan, Pervasive transcription: detecting functional RNAs in bacteria. Transcription 5, e944039 (2014).

30. M. C. Lybecker, D. S. Samuels, Small RNAs of *Borrelia burgdorferi*: characterizing functional regulators in a sea of sRNAs. Yale J Biol Med 90, 317–323 (2017).

31. D. Drecktrah, L. S. Hall, P. Rescheneder, M. Lybecker, D. S. Samuels, The Stringent Response-Regulated sRNA Transcriptome of *Borrelia burgdorferi*. Front Cell Infect Microbiol 8, 231 (2018).

32. N. Popitsch, I. Bilusic, P. Rescheneder, R. Schroeder, M. Lybecker, Temperature-dependent sRNA transcriptome of the Lyme disease spirochete. BMC Genomics 18, 28 (2017).

33. W. K. Arnold et al., RNA-Seq of *Borrelia burgdorferi* in Multiple Phases of Growth Reveals Insights into the Dynamics of Gene Expression, Transcriptome Architecture, and Noncoding RNAs. PLoS One 11, e0164165 (2016).

34. E. Petroni et al., Extensive diversity in RNA termination and regulation revealed by transcriptome mapping for the Lyme pathogen *Borrelia burgdorferi*. Nat Commun 14, 3931 (2023).

35. D. N. Medina-Pérez et al., The intergenic small non-coding RNA ittA is required for optimal infectivity and tissue tropism in *Borrelia burgdorferi*. PLoS Pathog 16, e1008423 (2020).

36. M. C. Lybecker, D. S. Samuels, Temperature-induced regulation of RpoS by a small RNA in *Borrelia burgdorferi*. Mol Microbiol 64, 1075–1089 (2007).

37. C. R. Savage et al., *Borrelia burgdorferi* SpoVG DNA- and RNA-Binding Protein Modulates the Physiology of the Lyme Disease Spirochete. J Bacteriol 200, (2018).

38. B. L. Jutras et al., Posttranscriptional self-regulation by the Lyme disease bacterium’s BpuR DNA/RNA-binding protein. J Bacteriol 195, 4915–4923 (2013).

39. T. C. Saylor et al., Quantitative analyses of interactions between SpoVG and RNA/DNA. Biochem Biophys Res Commun 654, 40–46 (2023).

40. S. Raghunandanan et al., A Fur family protein BosR is a novel RNA-binding protein that controls rpoS RNA stability in the Lyme disease pathogen. Nucleic Acids Res 52, 5320–5335 (2024).

41. B. L. Jutras et al., Bpur, the Lyme disease spirochete’s PUR domain protein: identification as a transcriptional modulator and characterization of nucleic acid interactions. J Biol Chem 288, 26220–26234 (2013).

42. A. C. Krusenstjerna, T. C. Saylor, W. K. Arnold, J. S. Tucker, B. Stevenson, *Borrelia burgdorferi* DnaA and the Nucleoid-Associated Protein EbfC Coordinate Expression of the. J Bacteriol 205, e0039622 (2023).

43. C. W. Sze et al., Carbon storage regulator A (CsrA(Bb)) is a repressor of *Borrelia burgdorferi* flagellin protein FlaB. Mol Microbiol 82, 851–864 (2011).

44. M. Zamba-Campero et al., Broadly conserved FlgV controls flagellar assembly and *Borrelia burgdorferi* dissemination in mice. Nat Commun 15, 10417 (2024).

45. M. C. Lybecker, C. A. Abel, A. L. Feig, D. S. Samuels, Identification and function of the RNA chaperone Hfq in the Lyme disease spirochete *Borrelia burgdorferi*. Mol Microbiol 78, 622–635 (2010).

46. T. Van Gundy et al., c-di-GMP regulates activity of the PlzA RNA chaperone from the Lyme disease spirochete. Mol Microbiol 119, 711–727 (2023).

47. D. L. Caly, P. W. O’Toole, S. A. Moore, The 2.2-Å structure of the HP0958 protein from Helicobacter pylori reveals a kinked anti-parallel coiled-coil hairpin domain and a highly conserved ZN-ribbon domain. J Mol Biol 403, 405–419 (2010).

48. H. Xu et al., *Borrelia burgdorferi* CheY2 Is Dispensable for Chemotaxis or Motility but Crucial for the Infectious Life Cycle of the Spirochete. Infect Immun 85, (2017).

49. M. A. Motaleb, S. Z. Sultan, M. R. Miller, C. Li, N. W. Charon, CheY3 of *Borrelia burgdorferi* is the key response regulator essential for chemotaxis and forms a long-lived phosphorylated intermediate. J Bacteriol 193, 3332–3341 (2011).

50. S. Stampfl, M. Doetsch, M. Beich-Frandsen, R. Schroeder, Characterization of the kinetics of RNA annealing and strand displacement activities of the *E. coli* DEAD-box helicase CsdA. RNA Biol 10, 149–156 (2013).

51. L. Rajkowitsch, R. Schroeder, Coupling RNA annealing and strand displacement: a FRET-based microplate reader assay for RNA chaperone activity. Biotechniques 43, 304, 306, 308 passim (2007).

52. M. Doetsch, T. Gstrein, R. Schroeder, B. Furtig, Mechanisms of StpA-mediated RNA remodeling. RNA Biol 7, 735–743 (2010).

53. M. Doetsch, B. Furtig, T. Gstrein, S. Stampfl, R. Schroeder, The RNA annealing mechanism of the HIV-1 Tat peptide: conversion of the RNA into an annealing-competent conformation. Nucleic Acids Res 39, 4405–4418 (2011).

54. M. Doetsch et al., Study of *E. coli* Hfq’s RNA annealing acceleration and duplex destabilization activities using substrates with different GC-contents. Nucleic Acids Res 41, 487–497 (2013).

55. B. G. Moreira, Y. You, R. Owczarzy, Cy3 and Cy5 dyes attached to oligonucleotide terminus stabilize DNA duplexes: predictive thermodynamic model. Biophys Chem 198, 36–44 (2015).

56. B. G. Moreira, Y. You, M. A. Behlke, R. Owczarzy, Effects of fluorescent dyes, quenchers, and dangling ends on DNA duplex stability. Biochem Biophys Res Commun 327, 473–484 (2005).

57. R. Landick, J. Stewart, D. N. Lee, Amino acid changes in conserved regions of the beta-subunit of *Escherichia coli* RNA polymerase alter transcription pausing and termination. Genes Dev 4, 1623–1636 (1990).

58. S. Phadtare, K. Severinov, RNA remodeling and gene regulation by cold shock proteins. RNA Biol 7, 788–795 (2010).

59. J. Li et al., The archaeal RNA chaperone TRAM0076 shapes the transcriptome and optimizes the growth of *Methanococcus maripaludis*. PLoS Genet 15, e1008328 (2019).

60. K. Nakaminami, D. T. Karlson, R. Imai, Functional conservation of cold shock domains in bacteria and higher plants. Proc Natl Acad Sci U S A 103, 10122–10127 (2006).

61. Z. Ouyang, R. K. Deka, M. V. Norgard, BosR (BB0647) controls the RpoN-RpoS regulatory pathway and virulence expression in *Borrelia burgdorferi* by a novel DNA-binding mechanism. PLoS Pathog 7, e1001272 (2011).

62. Z. Ouyang, J. Zhou, C. A. Brautigam, R. K. Deka, M. V. Norgard, Identification of a core sequence for the binding of BosR to the *rpoS* promoter region in *Borrelia burgdorferi*. Microbiology 161, 931 (2015).

63. C. Mason, X. Liu, S. Prabhudeva, Z. Ouyang, The CXXC Motifs Are Essential for the Function of BosR in *Borrelia burgdorferi*. Front Cell Infect Microbiol 9, 109 (2019).

64. J. A. Hyde, D. K. Shaw, R. Smith Iii, J. P. Trzeciakowski, J. T. Skare, The BosR regulatory protein of Borrelia burgdorferi interfaces with the RpoS regulatory pathway and modulates both the oxidative stress response and pathogenic properties of the Lyme disease spirochete. Mol Microbiol 74, 1344–1355 (2009).

65. Z. Ouyang et al., BosR (BB0647) governs virulence expression in *Borrelia burgdorferi*. Mol Microbiol 74, 1331–1343 (2009).

66. M. Modic, M. Adamek, J. Ule, The impact of IDR phosphorylation on the RNA binding profiles of proteins. Trends Genet 40, 580–586 (2024).

67. M. T. Lovci, M. H. Bengtson, K. B. Massirer, Post-Translational Modifications and RNA-Binding Proteins. Adv Exp Med Biol 907, 297–317 (2016).

68. D. S. M. Ottoz, L. E. Berchowitz, The role of disorder in RNA binding affinity and specificity. Open Biol 10, 200328 (2020).

69. G. Priyanka, E. J. Raj, N. P. Prabhu, Liquid-liquid phase separation of intrinsically disordered proteins: Effect of osmolytes and crowders. Prog Mol Biol Transl Sci 211, 249–269 (2025).

70. E. C. Wheeler, E. L. Van Nostrand, G. W. Yeo, Advances and challenges in the detection of transcriptome-wide protein-RNA interactions. Wiley Interdiscip Rev RNA 9, (2018).

71. D. S. Samuels, Electrotransformation of the spirochete *Borrelia burgdorferi*. Methods Mol Biol 47, 253–259 (1995).

72. D. S. Samuels, D. Drecktrah, L. S. Hall, Genetic Transformation and Complementation. Methods Mol Biol 1690, 183–200 (2018).

73. J. R. Izac et al., Development and optimization of OspC chimeritope vaccinogens for Lyme disease. Vaccine 38, 1915–1924 (2020).

74. D. Wagh et al., Borreliacidal activity of Borrelia metal transporter A (BmtA) binding small molecules by manganese transport inhibition. Drug Des Devel Ther 9, 805–816 (2015).

75. W. K. Boyle, H. N. Sorensen, T. J. Bourret, Essential Components of *Borreliella* (*Borrelia*) *burgdorferi In Vitro* Transcription Assays. J Vis Exp, (2022).

76. W. K. Boyle et al., Establishment of an *in vitro* RNA polymerase transcription system: a new tool to study transcriptional activation in *Borrelia burgdorferi*. Sci Rep 10, 8246 (2020).

