## Supplementary Information for "Identification of proteins exhibiting *in vitro* RNA chaperone activity through gradient profiling in the Lyme disease spirochete, *Borrelia burgdorferi*"

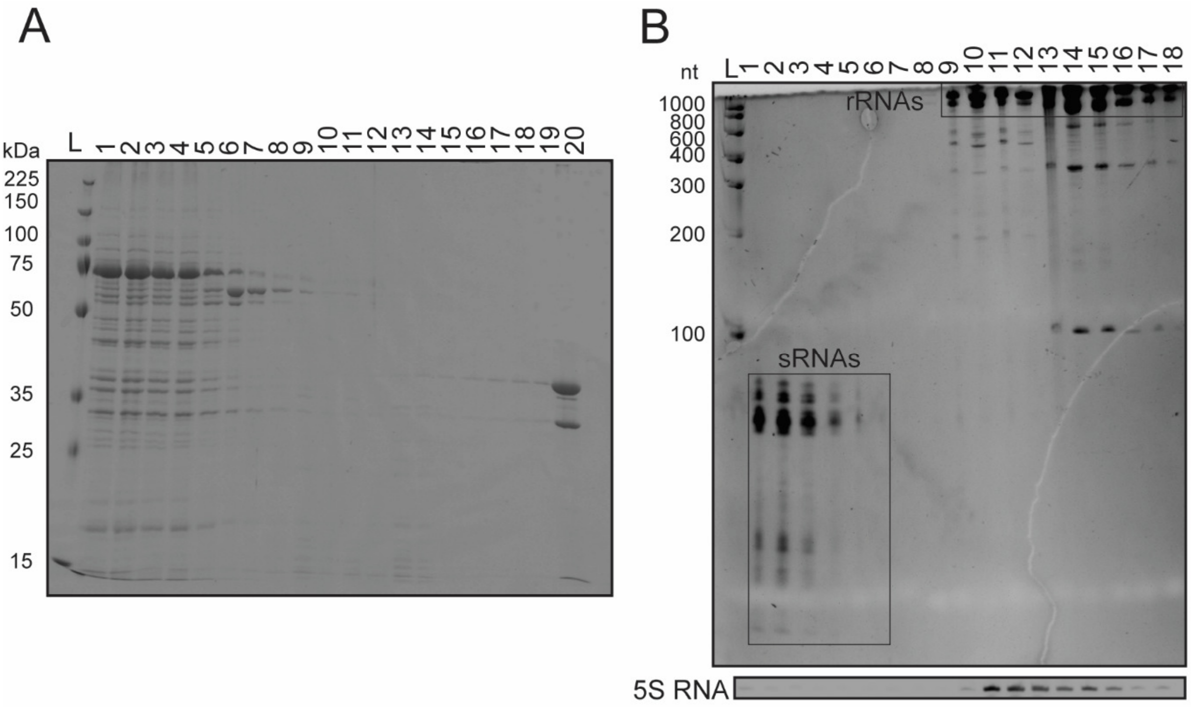


**Figure S1. Glycerol gradient fractionation validation.** (A) Coomassie Brilliant Blue-stained SDS-PAGE gel detecting proteins in the fractions from the glycerol gradient. The protein molecular weight ladder is denoted (L). (B) Ethidium bromide-stained TBE-UREA gel detecting RNA in fractions from the glycerol gradient. The RNA nucleotide ladder is denoted (L). Small RNAs and rRNAs are marked by black boxes. A 5S RNA northern blot validates the separation of the fractions.

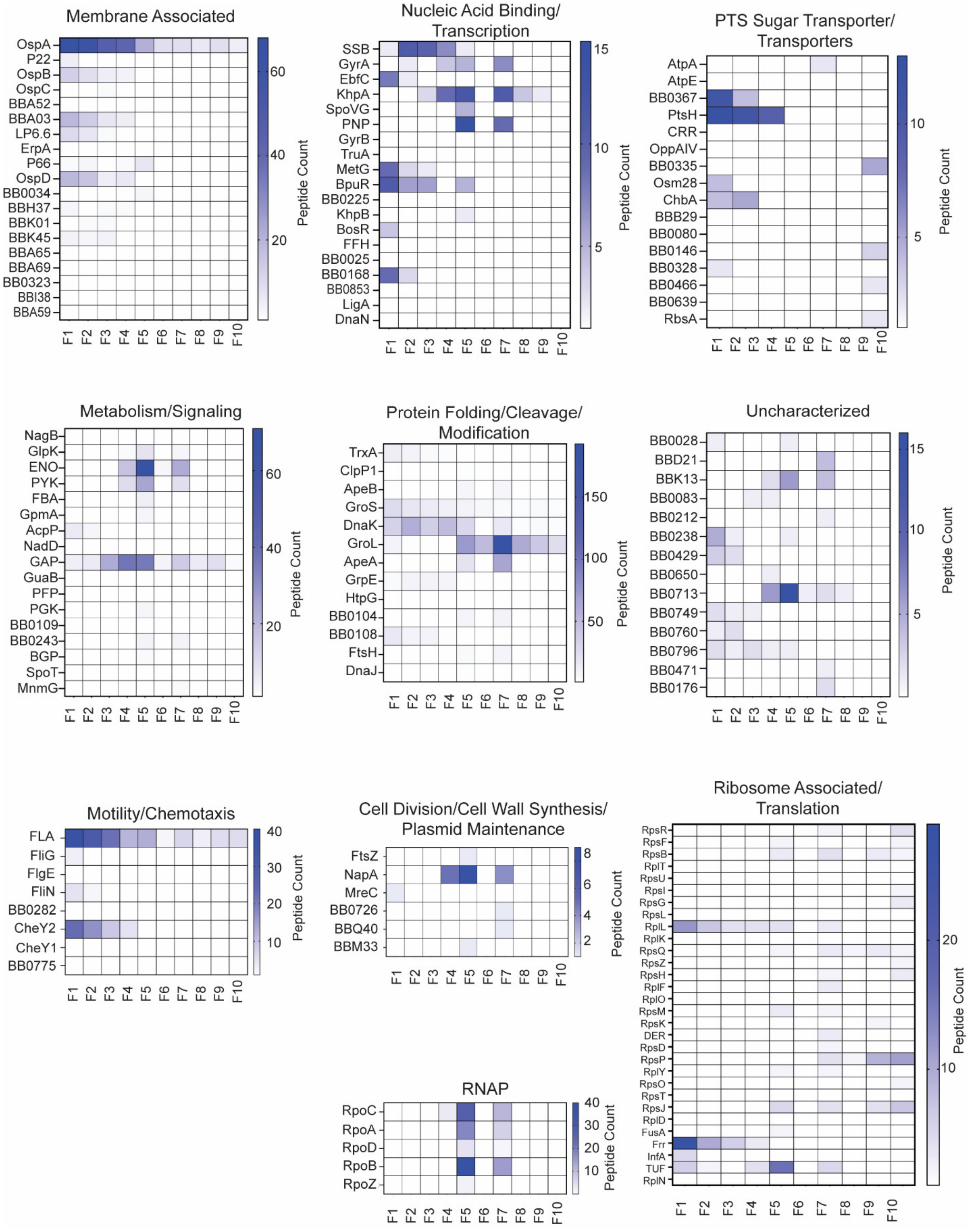
Membrane Associated
OspA
P22
OspB
OspC
BBA52
BBA03
LP6.6
ErpA
P66
OspD
BB0034
BBH37
BBK01
BBK45
BBA65
BBA69
BB0323
BBI38
BBA59
F1 F2 F3 F4 F5 F6 F7 F8 F9 F10
60
40
20
Peptide Count
Nucleic Acid Binding/ Transcription
SSB
GyrA
EbfC
KhpA
SpoVG
PNP
GyrB
TruA
MetG
BpuR
BB0225
KhpB
BosR
FFH
BB0025
BB0168
BB0853
LigA
DnaN
F1 F2 F3 F4 F5 F6 F7 F8 F9 F10
15
10
5
Peptide Count
PTS Sugar Transporter/ Transporters
AtpA
AtpE
BB0367
PtsH
CRR
OppAIV
BB0335
Osm28
ChbA
BBB29
BB0080
BB0146
BB0328
BB0466
BB0639
RbsA
F1 F2 F3 F4 F5 F6 F7 F8 F9 F10
10
5
Peptide Count
Metabolism/Signaling
NagB
GlpK
ENO
PYK
FBA
GpmA
AcpP
NadD
GAP
GuaB
PFP
PGK
BB0109
BB0243
BGP
SpoT
MnmG
F1 F2 F3 F4 F5 F6 F7 F8 F9 F10
60
40
20
Peptide Count
Protein Folding/Cleavage/ Modification
TrxA
ClpP1
ApeB
GroS
DnaK
GroL
ApeA
GrpE
HtpG
BB0104
BB0108
FtsH
DnaJ
F1 F2 F3 F4 F5 F6 F7 F8 F9 F10
150
100
50
Peptide Count
Uncharacterized
BB0028
BBD21
BBK13
BB0083
BB0212
BB0238
BB0429
BB0650
BB0713
BB0749
BB0760
BB0796
BB0471
BB0176
F1 F2 F3 F4 F5 F6 F7 F8 F9 F10
15
10
5
Peptide Count
Motility/Chemotaxis
FLA
FliG
FlgE
FliN
BB0282
CheY2
CheY1
BB0775
F1 F2 F3 F4 F5 F6 F7 F8 F9 F10
40
30
20
10
Peptide Count
Cell Division/Cell Wall Synthesis/ Plasmid Maintenance
FtsZ
NapA
MreC
BB0726
BBQ40
BBM33
F1 F2 F3 F4 F5 F6 F7 F8 F9 F10
8
6
4
2
Peptide Count
RNAP
RpoC
RpoA
RpoD
RpoB
RpoZ
F1 F2 F3 F4 F5 F6 F7 F8 F9 F10
40
30
20
10
Peptide Count
Ribosome Associated/ Translation
RpsR
RpsF
RpsB
RplT
RpsU
RpsI
RpsG
RpsL
RplL
RplK
RpsQ
RpsZ
RpsH
RplF
RplO
RpsM
RpsK
DER
RpsD
RpsP
RplY
RpsO
RpsT
RpsJ
RplD
FusA
Frr
InfA
TUF
RplN
F1 F2 F3 F4 F5 F6 F7 F8 F9 F10
20
10
Peptide Count

**Figure S2. Heat maps of in-gradient distributions of major groups of proteins identified in fractions 1-10 (F1-10).**

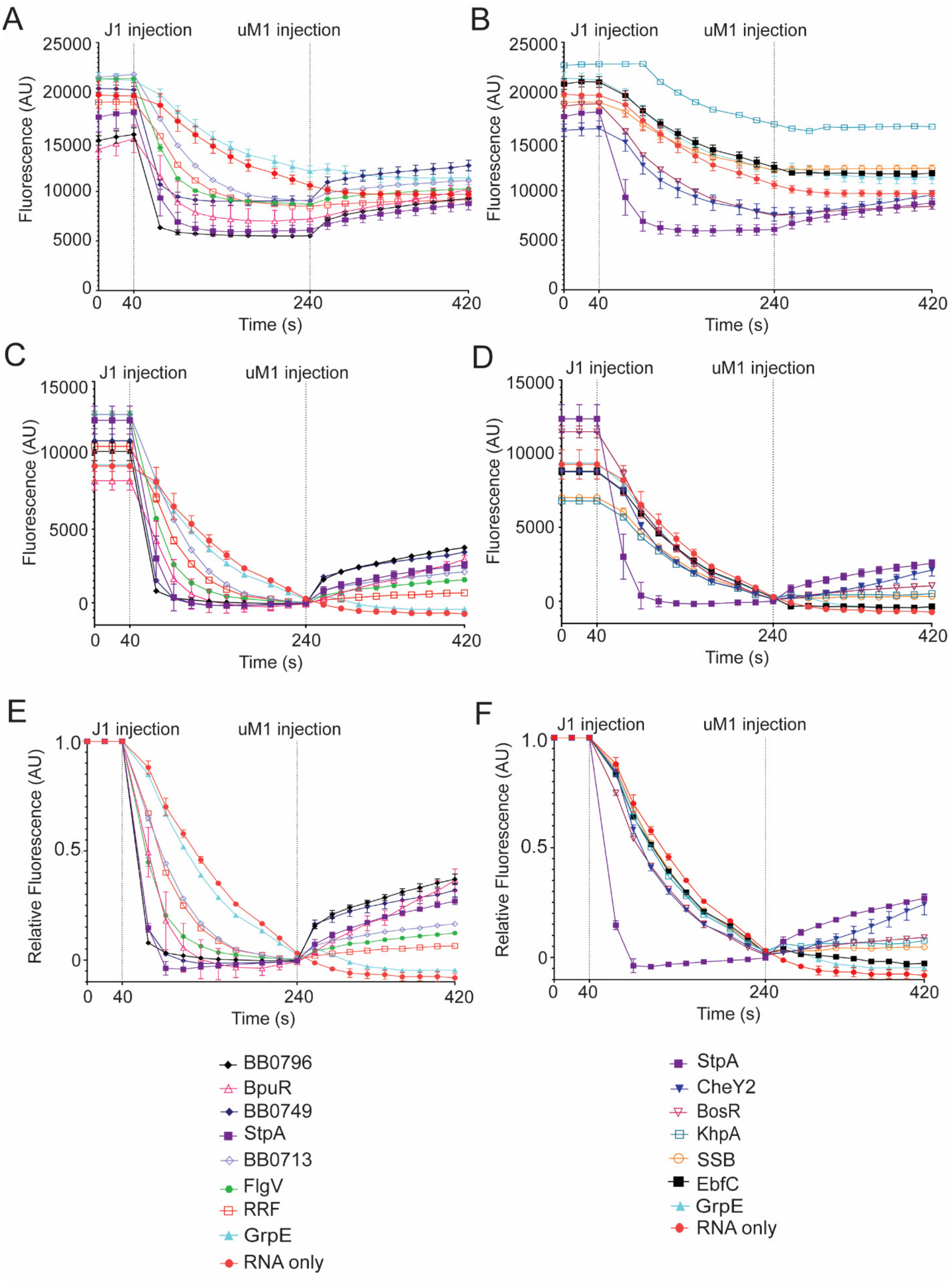
A
B
C
D
E
F
J1 injection
uM1 injection
Fluorescence (AU)
Relative Fluorescence (AU)
Time (s)
0
40
240
420
25000
20000
15000
10000
5000
1.0
0.5
BB0796
BpuR
BB0749
StpA
BB0713
FlgV
RRF
GrpE
RNA only
CheY2
BosR
KhpA
SSB
EbfC

**Figure S3**. **Normalization of in-solution annealing and strand displacement data**. (A and B) Relative fluorescent units of each time-resolved curve. (C and D) Normalization of time-resolved curves to 0 at 240s. (E and F) Normalization of time-resolved curves to 1 at 40s. Each vertical dotted line represents the time that RNA was injected into each sample, J1 injected at 40s and uM1 at 240s. RNA without protein (red circle) or GrpE (teal triangle) served as negative controls. StpA, an RNA chaperone from *E. coli*, was used as a positive control (purple square). *B. burgdorferi* proteins with annealing rates at least 2-fold higher than RNA only are displayed in panels A, C and E). Whereas *B. burgdorferi* proteins with annealing rates less than 2-fold compared to RNA only are displayed in panels B, D and F. Error bars represent the standard error of the mean (SEM) of three replicates. Error bars represent the standard error of the mean (SEM) of three replicates.

A

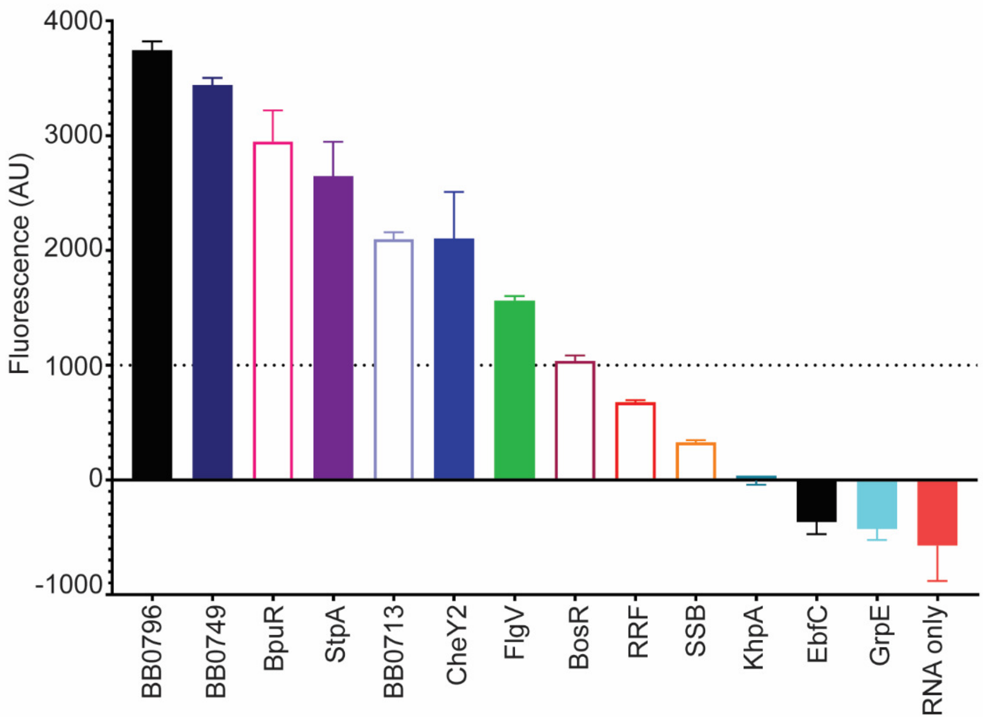


**Figure S4. Change in fluorescence during strand displacement reaction.** (A) The change in fluorescent units from the start (240 s) and the end (420 s) of the strand displacement reaction. Proteins are labeled on the x-axis. The dotted line indicates the positive strand displacement threshold value of 1000 AU. Error bars represent the standard error of the mean (SEM) of three replicates.

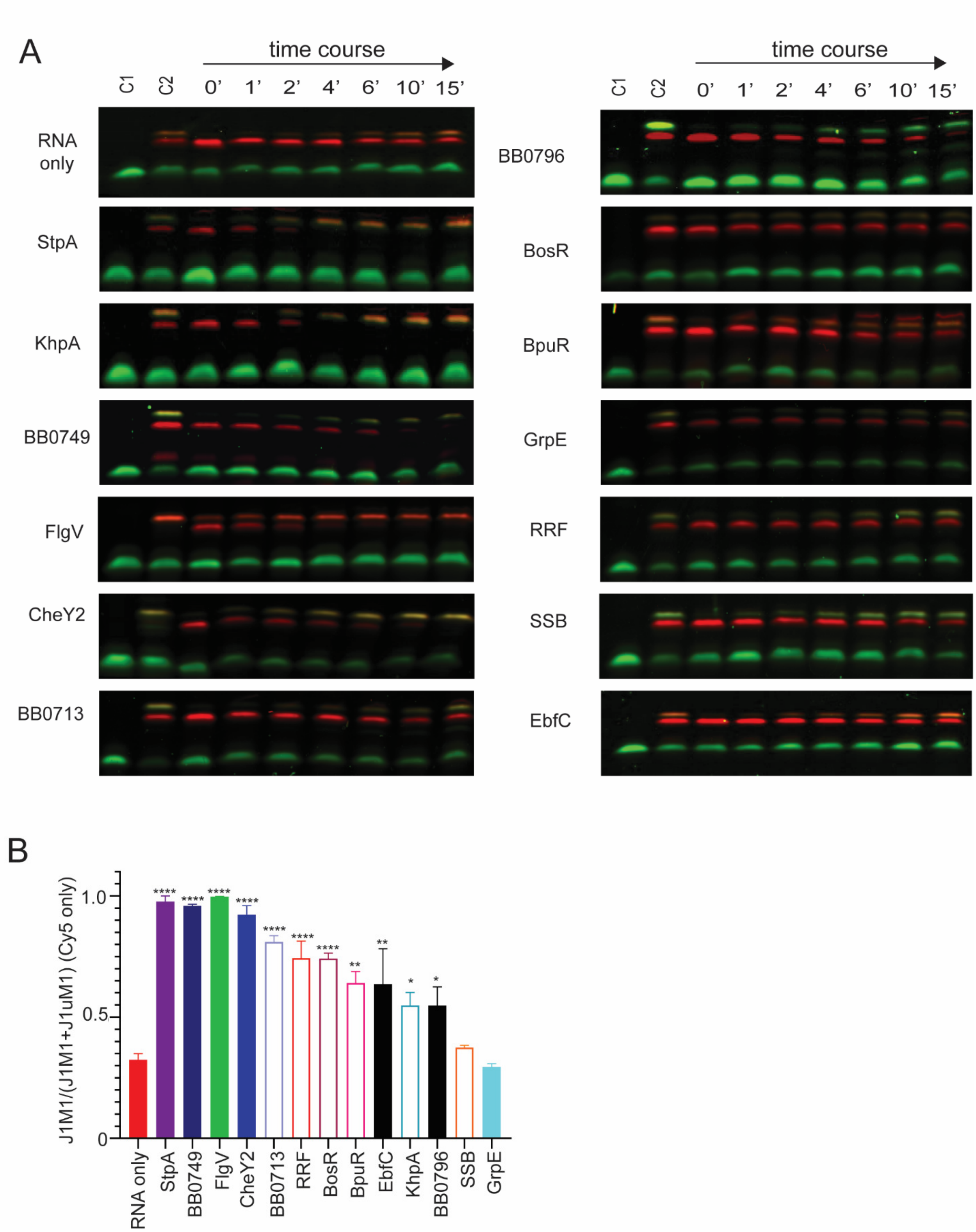


**Figure S5. Representative images of RNA annealing gels and end-point analysis.** (A) Representative images of RNA annealing gels. The gels shown are overlays of images scanned on an Azuer Sapphire fluorescence imager using the Cy3 and Cy5 channels. Control 1 (C1) is M1

RNA (green band) and control 2 (C2) is preformed J1M1 (yellow band) and uM1J1 (red band). (B) A bar graph illustrating the ratio of M1J1 dsRNA to total RNA at the endpoint of the RNA annealing assay. Annealing endpoint signals were quantified by ImageJ and analyzed by a two-way ANOVA with multiple comparisons (p-value $\leq$0.05, *, p-value $\leq$0.01, **, p-value $\leq$0.0001, ****). Bar colors correspond to symbol on gel-annealing graph (Fig 3). Error bars represent the SEM of three replicates.

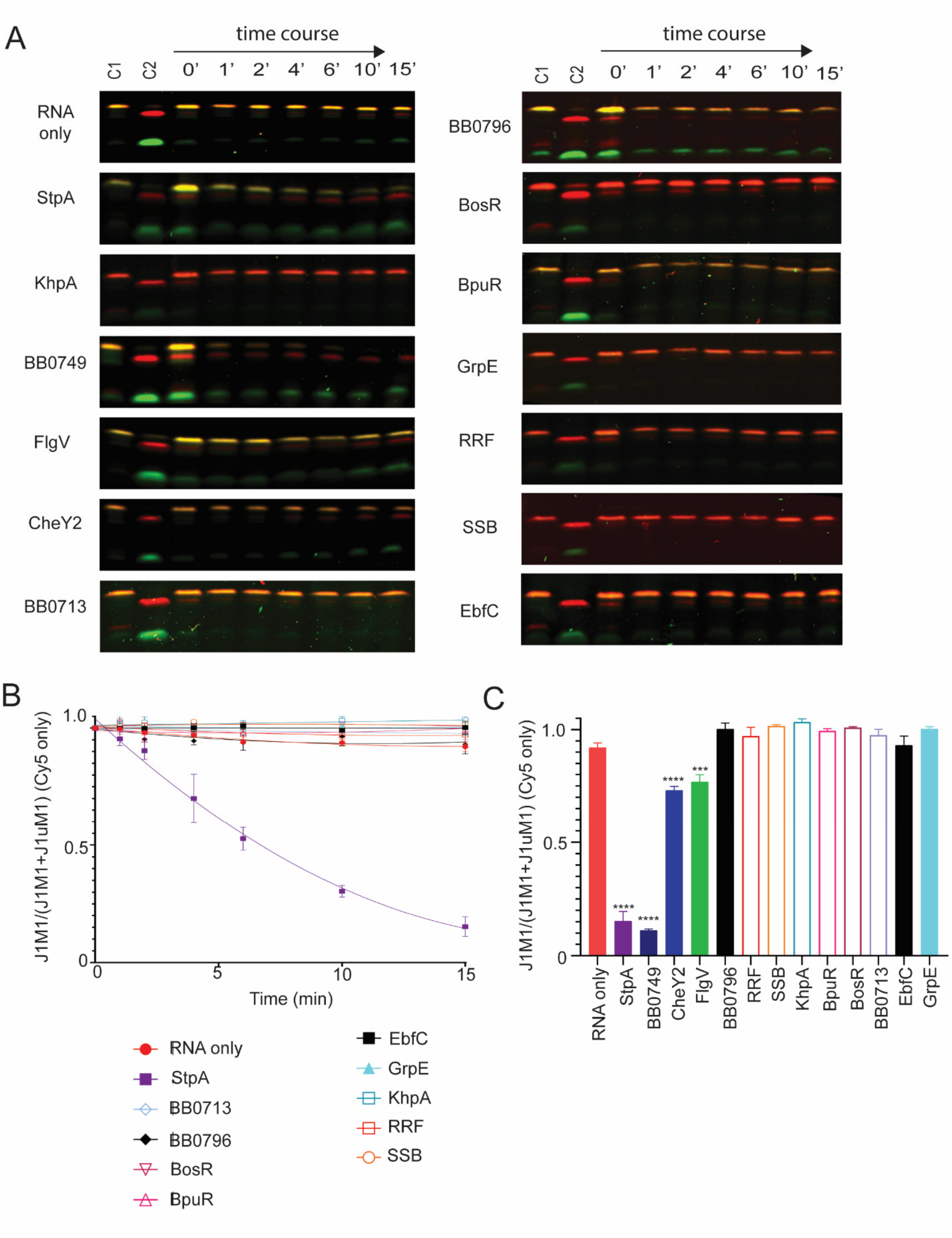


**Figure S6. Representative images of RNA strand displacement gels, displacement curves of proteins without stand displacement activity and end-point analyses.** (A) Representative images of RNA strand displacement gels. The gels shown are overlays of images scanned on an Azuer Sapphire fluorescence imager using the Cy3 and Cy5 channels. Control (C1) is preformed J1M1 RNA (yellow band) and control 2 (C2) is preformed J1uM1 (red band). The strand displacement reaction is initiated by adding unlabeled M1 (uM1) to preformed J1M1 in the presence or absence of protein. Samples are withdrawn from the reaction over time and analyzed by native polyacrylamide gel electrophoresis. The amount of preformed J1M1 dsRNA (yellow band) will decrease with a concomitant increase in the J1uM1 dsRNA (red band) formation over time if the protein has strand displacement activity. (B) Normalized non-linear regression of the ratio of J1M1 dsRNA to total RNA for each time point yielded stand displacement curves. Only the Cy5 channel was quantified and used for graphing the curves. RNA only (red circle) and GrpE (teal triangle) were used in the assay as negative controls, while StpA (purple square) was used as a positive control. BB0713 (open lavender diamond), RRF (open red square), BosR (open magenta upside-down triangle), BpuR (open pink triangle), EbfC (black square), KhpA (open teal square), BB0796 (black diamond) and SSB (open orange circle) did not display stand displacement activity. Error bars represent the SEM of three replicates. (C) A bar graph illustrating the ratio M1J1 dsRNA to total RNA at the endpoint of the RNA strand displacement assay. Strand displacement endpoint signals were quantified by Image J and analyzed by a two-way ANOVA with multiple comparisons (p-value $\leq$0.001, ***, p-value $\leq$0.0001, ****). Bar graph colors correspond to symbol on gel-annealing graph (Fig 4). Error bars represent the SEM of three replicates.

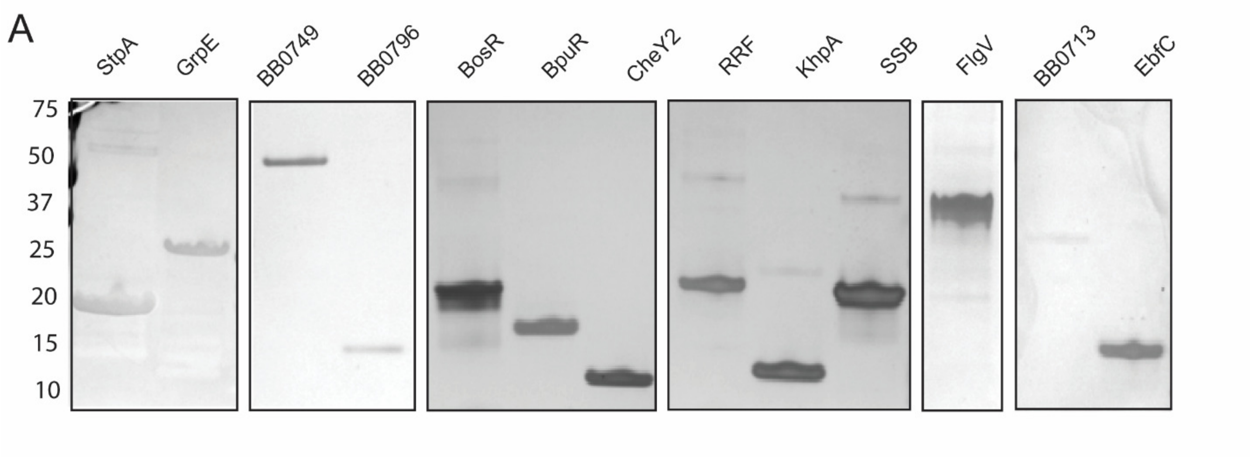


**Figure S7. Recombinant Proteins were resolved on precast Criterion Any kD™ TGX Stain-Free™ Protein Gels (BioRad) and visualized by silver staining to assess protein purity.**

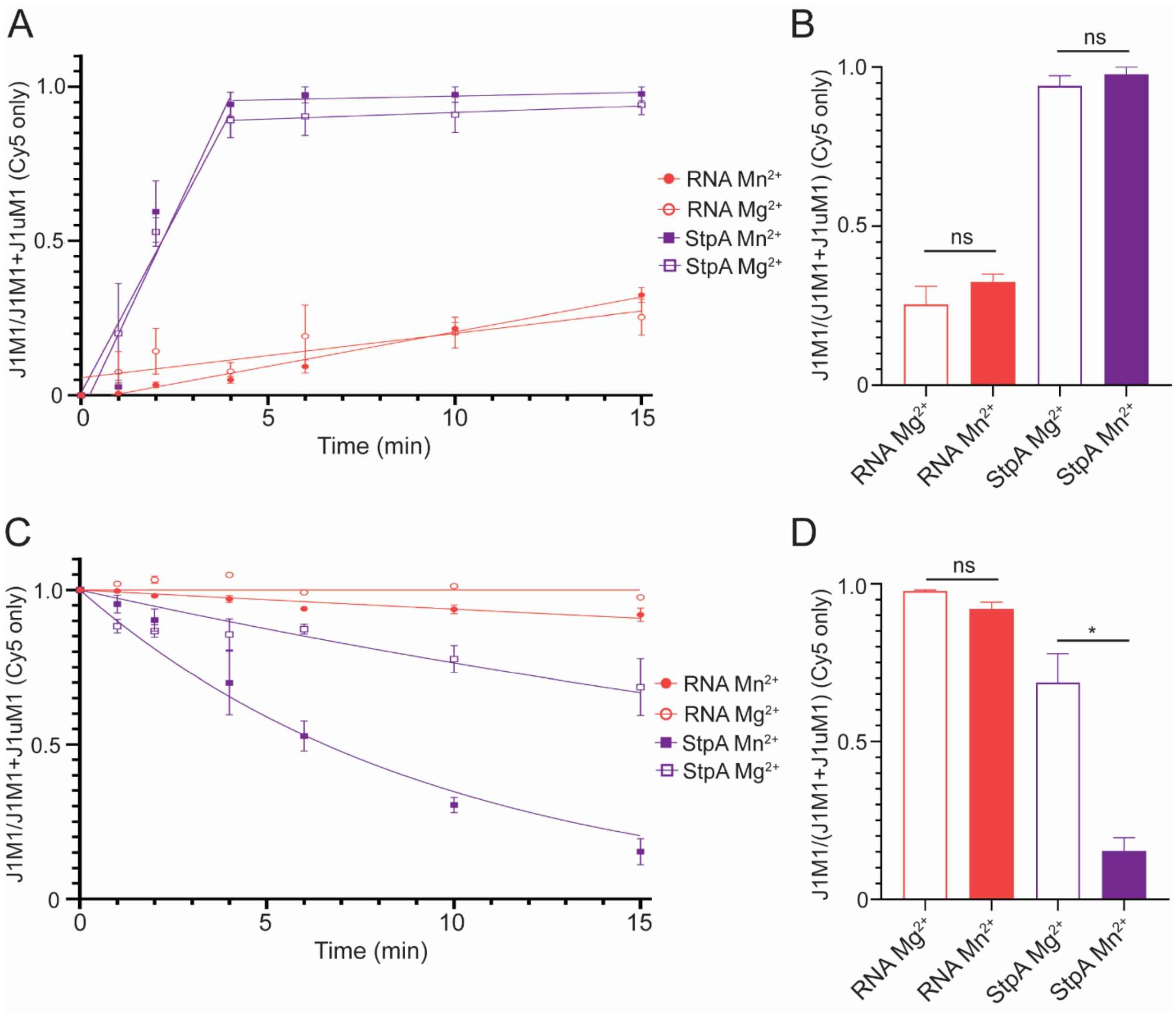


**Figure S8. StpA displays stronger strand displacement activity in buffer with $Mn^{2+}$ compared to $Mg^{2+}$ in gel assays.** (A) RNA annealing curves were generated from normalized non-linear regression of the ratio of J1M1 dsRNA to total RNA for each time point for samples: RNA only $Mg^{2+}$ (open red circle), RNA only $Mn^{2+}$ (red circle), StpA $Mg^{2+}$ (open purple square), StpA $Mn^{2+}$ (purple square). (B) A bar graph illustrating the ratio of M1J1 dsRNA to total RNA at the endpoint of the RNA annealing assay. Annealing endpoint signals were quantified by ImageJ and analyzed by a Welch's t-Test (p-value >0.05, ns). (C) RNA strand displacement curves were generated from normalized non-linear regression of the ratio of J1M1 dsRNA to total RNA for each time point for samples: RNA only $Mg^{2+}$ (open red circle), RNA only $Mn^{2+}$ (red circle), StpA $Mg^{2+}$ (open purple square), StpA $Mn^{2+}$ (purple square). (D) A bar graph illustrating the ratio of M1J1 dsRNA to total RNA at the endpoint of the RNA strand displacement assay. Strand displacement endpoint signals were quantified by Image J and analyzed by a Welch's t-Test (p-value >0.05, ns, p-value $\leq$0.05, *). Bar graph colors correspond to symbol on gel-annealing graph (Fig 4). Error bars represent the SEM of three replicates.

Table S2: Alphafold3 strucutres, protein descriptions, and protein references

| Alphafold Prediction | Description | References |
|---|---|---|
| 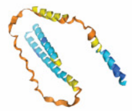 | FlgV; Cell membrane; Multi-pass membrane protein; Transmembrane domain | Lybecker *et al.* 2010<br>Petroni *et al.* 2023<br>Zamba-Campero *et al.* 2024 |
| 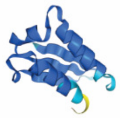 | KhpA; RNA binding protein; KH-domain | Olejniczak *et al.* 2022 |
| 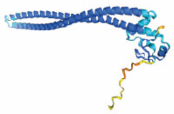 | BB0713; C4-type zinc ribbon domain-containing protein | Mueller *et al.* 2006<br>Kariu *et al.* 2013 |
| 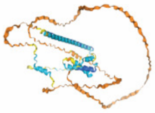 | BB0749; IDR, Unknown function | Miller *et al.* 2013<br>Iyer *et al.* 2015<br>Jutras *et al.* 2012<br>Mongodin *et al.* 2013 |
| 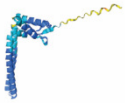 | BB0796; coiled coil and signal domain; Skp homolog | Schulze *et al.* 2010<br>Jutras *et al.* 2019 |
| 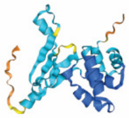 | BosR; oxidative stress regulator; belongs to the FUR family; CXXC binding domain | Boylan *et al.* 2003<br>Ouyang *et al.* 2011<br>Mason *et al.* 2019<br>Grassmann *et al.* 2023 |
| 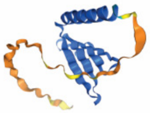 | BpuR; PUR domain | Jutras *et al.* 2013<br>Jutras *et al.* 2019 |
| 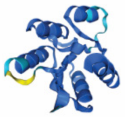 | CheY2; two component response regulator; CheY-1 like superfamily | Li *et al.* 2002<br>Motaleb *et al.* 2011<br>Xu *et al.* 2016 |
| 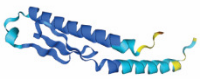 | EbfC; nucleoid-associat-ed protein | Babb *et al.* 2006<br>Riley *et al.* 2009<br>Krusenstjerna *et al.* 2023 |
| 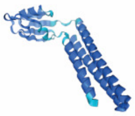 | RRF; Ribosome recycling factor | Saylor *et al.* 2022 |
| 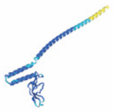 | GrpE; Head domain nucleotide exchange factor, coiled-coil domain | Tilly *et al.* 1993 |
| 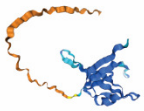 | SSB; nucleic acid single stranded binding | Chenail *et al.* 2012<br>Huang *et al.* 2017 |

Very High (pIDDT > 90) Confident (90 < pIDDT > 70) Low (70 > pIDDT > 50) Very Low (pIDDT < 50) (*1-28*)

| Proteins | Annealing and Strand Displacement | | | | Unwinding and Anti-termination | |
|---|---|---|---|---|---|---|
| | In-solution | | Gel-based | | *E. coli* | *in vitro* |
| | A | SD | A | SD | | |
| StpA | + | + | + | + | + | + |
| BB0796 | + | + | + | - | + | + |
| BpuR | + | + | + | - | + | + |
| BB0749 | + | + | + | + | - | + |
| BB0713 | + | + | + | - | + | + |
| FlgV | + | + | + | + | + | + |
| RRF | + | - | + | - | + | - |
| CheY2 | + | + | + | + | + | - |
| BosR | + | + | + | - | + | - |
| KhpA | - | - | + | - | - | + |
| SSB | - | - | - | - | N/A | - |
| EbfC | - | - | + | - | + | - |
| GrpE | - | - | - | - | - | - |
| RNA only | - | - | - | - | - | - |

**Table S5: Summary of RNA chaperone activity of each protein.** Annealing and strand displacement assays are denoted as A and SD respectively. The *in vitro* unwinding assay is denoted as *in vitro* and the antitermination assay in *E. coli* is denoted as *E. coli*. A (+) denotes that a protein demonstrated that RNA chaperone activity, while a (-) denotes proteins without that activity. N/A denotes an assay that was not performed.

### Supplementary References


1. K. Babb *et al.*, *Borrelia burgdorferi* EbfC, a novel, chromosomally encoded protein, binds specific DNA sequences adjacent to *erp* loci on the spirochete's resident cp32 prophages. *J Bacteriol* **188**, 4331-4339 (2006).
2. J. A. Boylan, J. E. Posey, F. C. Gherardini, Borrelia oxidative stress response regulator, BosR: a distinctive Zn-dependent transcriptional activator. *Proc Natl Acad Sci U S A* **100**, 11684-11689 (2003).
3. A. M. Chenail *et al.*, *Borrelia burgdorferi* cp32 BpaB modulates expression of the prophage NucP nuclease and SsbP single-stranded DNA-binding protein. *J Bacteriol* **194**, 4570-4578 (2012).
4. A. A. Grassmann *et al.*, BosR and PlzA reciprocally regulate RpoS function to sustain *Borrelia burgdorferi* in ticks and mammals. *J Clin Invest* **133**, (2023).
5. S. H. Huang, M. R. Cozart, M. A. Hart, K. Kobryn, The *Borrelia burgdorferi* telomere resolvase, ResT, possesses ATP-dependent DNA unwinding activity. *Nucleic Acids Res* **45**, 1319-1329 (2017).
6. R. Iyer *et al.*, Stage-specific global alterations in the transcriptomes of Lyme disease spirochetes during tick feeding and following mammalian host adaptation. *Mol Microbiol* **95**, 509-538 (2015).
7. B. L. Jutras *et al.*, EbfC (YbaB) is a new type of bacterial nucleoid-associated protein and a global regulator of gene expression in the Lyme disease spirochete. *J Bacteriol* **194**, 3395-3406 (2012).
8. B. L. Jutras *et al.*, Bpur, the Lyme disease spirochete's PUR domain protein: identification as a transcriptional modulator and characterization of nucleic acid interactions. *J Biol Chem* **288**, 26220-26234 (2013).
9. B. L. Jutras *et al.*, Posttranscriptional self-regulation by the Lyme disease bacterium's BpuR DNA/RNA-binding protein. *J Bacteriol* **195**, 4915-4923 (2013).
10. B. L. Jutras *et al.*, The Lyme disease spirochete's BpuR DNA/RNA-binding protein is differentially expressed during the mammal-tick infectious cycle, which affects translation of the SodA superoxide dismutase. *Mol Microbiol* **112**, 973-991 (2019).
11. T. Kariu, X. Yang, C. B. Marks, X. Zhang, U. Pal, Proteolysis of BB0323 results in two polypeptides that impact physiologic and infectious phenotypes in *Borrelia burgdorferi*. *Mol Microbiol* **88**, 510-522 (2013).
12. A. C. Krusenstjerna, T. C. Saylor, W. K. Arnold, J. S. Tucker, B. Stevenson, *Borrelia burgdorferi* DnaA and the Nucleoid-Associated Protein EbfC Coordinate Expression of the. *J Bacteriol* **205**, e0039622 (2023).
13. C. Li *et al.*, Asymmetrical flagellar rotation in *Borrelia burgdorferi* nonchemotactic mutants. *Proc Natl Acad Sci U S A* **99**, 6169-6174 (2002).
14. M. C. Lybecker, C. A. Abel, A. L. Feig, D. S. Samuels, Identification and function of the RNA chaperone Hfq in the Lyme disease spirochete Borrelia burgdorferi. *Mol Microbiol* **78**, 622-635 (2010).
15. C. Mason, X. Liu, S. Prabhudeva, Z. Ouyang, The CXXC Motifs Are Essential for the Function of BosR in *Borrelia burgdorferi*. *Front Cell Infect Microbiol* **9**, 109 (2019).
16. C. L. Miller, S. L. Karna, J. Seshu, *Borrelia* host adaptation Regulator (BadR) regulates *rpoS* to modulate host adaptation and virulence factors in *Borrelia burgdorferi*. *Mol Microbiol* **88**, 105-124 (2013).

17. E. F. Mongodin *et al.*, Inter- and intra-specific pan-genomes of *Borrelia burgdorferi sensu lato*: genome stability and adaptive radiation. *BMC Genomics* **14**, 693 (2013).
18. M. A. Motaleb, S. Z. Sultan, M. R. Miller, C. Li, N. W. Charon, CheY3 of *Borrelia burgdorferi* is the key response regulator essential for chemotaxis and forms a long-lived phosphorylated intermediate. *J Bacteriol* **193**, 3332-3341 (2011).
19. M. Mueller *et al.*, Identification of *Borrelia burgdorferi* ribosomal protein L25 by the phage surface display method and evaluation of the protein's value for serodiagnosis. *J Clin Microbiol* **44**, 3778-3780 (2006).
20. M. Olejniczak, X. Jiang, M. M. Basczok, G. Storz, KH domain proteins: Another family of bacterial RNA matchmakers? *Mol Microbiol* **117**, 10-19 (2022).
21. Z. Ouyang, R. K. Deka, M. V. Norgard, BosR (BB0647) controls the RpoN-RpoS regulatory pathway and virulence expression in *Borrelia burgdorferi* by a novel DNA-binding mechanism. *PLoS Pathog* **7**, e1001272 (2011).
22. E. Petroni *et al.*, Extensive diversity in RNA termination and regulation revealed by transcriptome mapping for the Lyme pathogen *Borrelia burgdorferi*. *Nat Commun* **14**, 3931 (2023).
23. S. P. Riley *et al.*, *Borrelia burgdorferi* EbfC defines a newly-identified, widespread family of bacterial DNA-binding proteins. *Nucleic Acids Res* **37**, 1973-1983 (2009).
24. T. C. Saylor *et al.*, *Borrelia burgdorferi*, the Lyme disease spirochete, possesses genetically-encoded responses to doxycycline, but not to amoxicillin. *PLoS One* **17**, e0274125 (2022).
25. R. J. Schulze, S. Chen, O. S. Kumru, W. R. Zuckert, Translocation of *Borrelia burgdorferi* surface lipoprotein OspA through the outer membrane requires an unfolded conformation and can initiate at the C-terminus. *Mol Microbiol* **76**, 1266-1278 (2010).
26. K. Tilly, J. Campbell, A *Borrelia burgdorferi* homolog of the *Escherichia coli rho* gene. *Nucleic Acids Res* **21**, 1040 (1993).
27. H. Xu *et al.*, *Borrelia burgdorferi* CheY2 Is Dispensable for Chemotaxis or Motility but Crucial for the Infectious Life Cycle of the Spirochete. *Infect Immun* **85**, (2017).
28. M. Zamba-Campero *et al.*, Broadly conserved FlgV controls flagellar assembly and *Borrelia burgdorferi* dissemination in mice. *Nat Commun* **15**, 10417 (2024).
